# Host-structured adult butterfly microbiomes share a common core across continents

**DOI:** 10.64898/2026.02.27.708436

**Authors:** Arne Weinhold, Andrea Pinos, Yenny Correa-Carmona, Kim L. Holzmann, Pedro Alonso-Alonso, Felipe Yon, Ingolf Steffan-Dewenter, Marcell K. Peters, Gunnar Brehm, Alexander Keller

## Abstract

The assembly of host-specific microbiota is critical for health and functioning of many insect pollinators. While social pollinators maintain core microbiota through social transmission, the factors driving microbiota assembly in solitary pollinators remain poorly understood. Lepidoptera are an important group of pollinators, but microbiome research has largely focused on their larval stage, while nectar-feeding adults have been widely ignored. Field-based studies of the adult butterfly microbiota are rare and geographically and taxonomically restricted.

Here, we characterize the microbiota of adult butterflies from natural environments along elevational and seasonal gradients across two continents spanning the Neotropical and Palearctic realms. Microbiota diversity and composition were primarily explained by host taxonomic identity, whereas geographic location and temperature had little effect. We found high microbial titers and a common core microbiota across seven butterfly subfamilies from temperate and tropical regions. A comparison with sympatric wild bees revealed partial taxonomic overlap but marked distinctions in lactic acid bacteria between Lepidoptera and Hymenoptera.

Together, our results demonstrate that host taxonomic identity, rather than environmental drivers, is the dominant force structuring the microbiota of adult butterflies. This highlights that the host-filtering capacity of solitary species has been largely underestimated, challenging previous assumptions that the microbiota of this important pollinator group is primarily environmentally driven and of limited functional significance.

## 2 Introduction

Microbes play an important role in pollination, acting as a third interaction partner between flowers and pollinating insects, providing benefits for both sides^1,2^. Social pollinators are well known for their ability to maintain microbes via conspecific interactions, resulting in conserved and host-specific core microbiota^3,4^. In contrast, solitary pollinators account for most pollinating species worldwide and their microbiota is assumed to be mainly shaped by environmental factors, as they acquire their microbiota primarily from environmental sources via plant-insect interaction networks^5^. Microbial sharing between flowers and insects has long been hypothesized^6,7^ and experimentally confirmed in solitary bees and butterflies^8,9^. Despite this, microbiota research has largely focused on hymenopteran pollinators^10^, while other insect orders (e.g. Hemiptera, Diptera or Lepidoptera) have received comparatively little attention^11,12^. Comparing pollinator microbiota across insect orders co-occurring in the same habitat enables a better evaluation of host-specific enrichment.

Lepidoptera are the second-largest insect order in terms of species number and play a crucial ecological role as important non-bee pollinators^13,14^. Despite a long research tradition, studies of the lepidopteran microbiota have focused almost exclusively on the phytophagous larval stage (caterpillars)^15,16^, while the adult stage (imago) has been widely ignored^17–20^. This bias was further amplified by laboratory assays using moth pest species, where adults had no access to flowers or nectar microbes, but were fed sterile sugar solutions^21,22^ or sampled immediately after emergence without prior feeding^23^. These studies on larvae and captive adults reinforced the widespread perception that Lepidoptera lack specific microbial associations^16,19,24^. However, adult butterflies differ markedly in gut morphology, lacking the simple tube-like guts of larvae but contain folds and villi-like structures that provide surfaces for microbial colonization^25^. Unlike larvae, adult guts are less alkaline, with more acidic pH conditions in the hindgut^26^ and digestive enzyme activity optimal at pH 6.4^27^. When directly compared, adults of true butterflies (Papilionoidea) harbor microbiota compositions clearly distinct from those of larvae^28–30^. That adults sampled from natural environments show contrasting microbiota compositions and substantial microbial loads has only recently been recognized in lepidopteran microbiome research^31^.

Nevertheless, comprehensive studies of the adult butterfly microbiota from natural environments remain scarce and are largely limited to Heliconiinae (Nymphalidae) from Central and South America^28,32–35^. These studies revealed high microbial titers in Neotropical butterflies and communities clearly distinct from those of food sources or trap baits^32,35^. Microbiota compositions were best explained by host taxonomic placement, rather than diet, contrasting with initial expectations that feeding preferences—nectar versus fruit, or pollen versus non-pollen—would be primary drivers^32,33^. Neotropical butterflies were consistently associated with a characteristic set of microbial taxa, dominated by insect-specific groups (*Orbus, Wolbachia, Spiroplasma*), lactic acid bacteria (*Enterococcus*, *Lactococcus*) and acetic acid bacteria (*Asaia*, *Commensalibacter*)^28,32–35^. Whether this pattern is specific to Neotropical species or a general signature of the adult butterfly microbiota remains unclear, as systematic samplings across distant climate zones are still lacking. Defining a ‘common core’ microbiota across different host families or locations is a crucial first step to distinguish potentially host-enriched taxa from transient microbes^3,36^. To date, global investigations have focused primarily on vertically transmitted endosymbionts such as *Wolbachia*^37,38^, and the presence of other microbial taxa must be inferred from reports of individual bacterial isolates^39,40^ or from non-host reads mined from genomic datasets^41^.

Here, we address this knowledge gap by systematically investigating the microbiota of adult butterflies across distinct geographic regions and climate zones, comparing species from the Neotropical and Palearctic realms. Samples were collected along an elevational gradient in Peru (subfamilies: Dismorphiinae, Pierinae, Coliadinae, Danainae and Heliconiinae), spanning largely undisturbed habitats from the species-rich lowland Amazon basin to montane forests, and in Germany (subfamilies: Dismorphiinae, Pierinae, Coliadinae Satyrinae and Nymphalinae), through repeated seasonal sampling from suburban areas. This design allows us to use temperature as environmental predictor of microbiota variation along a gradient ranging from 6.7°C to 26.1°C, to disentangle environmental influences from host filtering effects. For broader host context, we compare butterfly microbiota to that of sympatric wild bees from a previous study sampled along the same Peruvian gradient^42^.

Expanding upon previous work centered on Heliconiinae, our sampling covers a broader taxonomic range with approximately 70% of samples belonging to Pierinae and Coliadinae. These ‘whites’ and ‘sulfurs’ are evolutionary model groups to study diversification dynamics in response to environmental temperatures^43^. Their widespread distribution across both study regions enabled us to assess whether environmental temperature, geographic location, or host taxonomy are the primary drivers of the adult butterfly microbiota. Within this context, we test the following hypotheses:

1. Environmental temperature shapes microbiota diversity in adult butterflies.
2. Microbial community composition is structured by host taxonomic identity.
3. Adult butterflies harbor a common core microbiota independent of geographic location.

We further compare common core taxa of adult butterflies with those reported from hymenopteran pollinators sampled from the same area in Peru^42^, highlighting distinctions among bee- and butterfly-associated groups. Together, these results offer a comprehensive characterization of the adult butterfly microbiota, including the first broad microbiome survey of European populations. This lays a foundation to further explore the ecological relevance and functional roles of adult-stage microbiota in butterflies.

## 3 Results

We compared the microbiota of adult butterflies across two distant countries (Peru and Germany) representing the Neotropics and the Palearctic realms (Figure 1A) within seven butterfly subfamilies: Dismorphiinae (n = 4), Pierinae (n = 73) and Coliadinae (n = 41) from the Pieridae family, and Danainae (n = 3), Satyrinae (n = 10), Heliconiinae (n = 27) and Nymphalinae (n = 9) belonging to the Nymphalidae family.

**Figure 1.**
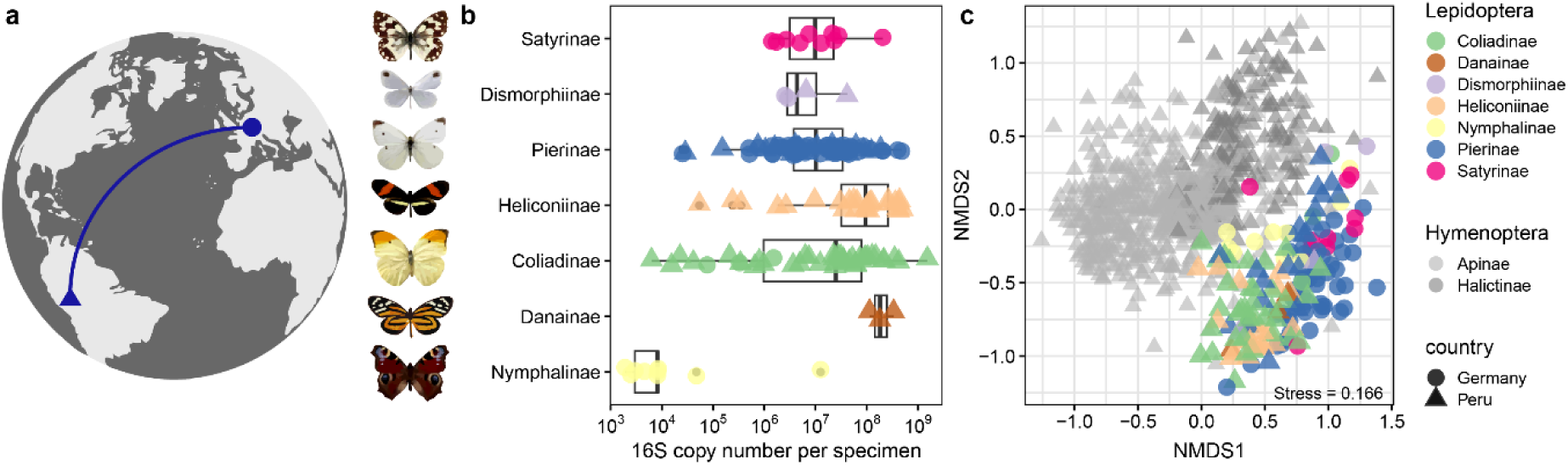
Adult butterflies harbor abundant microbiomes with host-structured community composition across continents. (**a**) Overview of adult butterfly sampling locations in Peru and Germany. Representative species are shown for each butterfly subfamily. (**b**) Absolute quantification of the bacterial load by 16S rRNA gene qPCR. (**c**) NMDS ordination based on Bray–Curtis dissimilarities comparing microbiome composition of Lepidoptera and Hymenoptera. Adult butterflies from Peru (n = 90) cluster together with butterflies from Germany (n = 77), while their microbiomes are distinct from those of sympatric wild bees from Peru (Apinae n = 670 and Halictinae n = 204).

### 3.1 Adult butterflies host abundant microbiota distinct from Hymenoptera

Quantitative analysis by 16S qPCR showed that the bacterial load of the different butterfly subfamilies spanned a broad range between 10^3^–10^9^ 16S copy numbers per specimen (Figure 1B). Adult butterflies harbored a median bacterial load around 1.85 × 10^7^ 16S copies (interquartile range: 2.6 × 10^6^ – 6.1 × 10^7^) which was strongly predicted by host genus identity in a linear model (adj. R² = 0.51, F_23,143_ = 8.6, *p* < 10^-15^) (Supplementary Figure S1A). A notable exception in abundance were *Aglais* (Nymphalinae), which were collected very early in season at low temperatures. Within individual subfamilies the bacterial load did not differ between countries (Supplementary Figure S1B). Using Type II ANOVA on a linear model showed a strong effect of host subfamily on bacterial load (partial R² = 0.30, *F*_6,159_ = 11.32, *p* < 10^-9^), whereas country had no significant effect (partial R² = 0.003, F_1,159_ = 0.49, p = 0.48).

For a broader insect order comparison, we combined the butterfly microbiota with that of wild bees, collected from the same locations in Peru^42^. Butterflies from Peru grouped more closely with those sampled from Germany, than with sympatric bees (Figure 1C). Overall microbial community composition differed between Hymenoptera and Lepidoptera, with host order explaining 8.2% of the variation in community composition (PERMANOVA, R^2^ = 0.082, F_1,1037_ = 92.1, P < 0.001).

Within adult butterflies, microbiota diversity was better explained by host taxonomic placement than by sampling location (Supplementary Figure S1C). A linear model including host subfamily and location explained 31% of variation in Shannon diversity (adj. R² = 0.31, F_9,157_ = 9.4, *p* < 10^-11^). We found a strong effect of host subfamily (Type II ANOVA partial R² = 0.17, *F*_6,157_ = 5.5, *p* < 0.0001), whereas sampling location had no significant effect (partial R² = 0.04, F_3,157_ = 2.1, p = 0.11). A similar result was obtained with a model including host tribe and country as predictor variables (Supplementary Figure S1C). As the sampling was not equal across countries, we additionally tested the effect of country on Shannon diversity within individual butterfly subfamilies (Supplementary Figure S1D). Still, country had no significant effect on microbial alpha diversity for Coliadinae (*p* = 0.12) or Pierinae (*p* = 0.62), while inferences are limited for Dismorphiinae (*p* = 0.33) due to the small sample size (Supplementary Figure S1D). Although samples had been collected from different continents, sampling location had minor influence on Shannon diversity compared to host taxonomic identity.

### 3.2 Influence of environmental gradients on adult butterfly microbiota diversity

Butterflies from Peru came from an elevational gradient, which showed strong linear correlation with environmental temperature (adj. R^2^ = 0.91). A linear model with only the Peru samples including host subfamily and temperature as predictors explained 26% of the variation in Shannon diversity (adj. R^2^ = 0.26, *F*_5,84_ = 7.2, *p* < 10^-4^). While the main effect was accounted to host subfamily (partial R² = 0.14, *F*_4,84_ = 3.5, *p* = 0.011), temperature showed a lower effect size but was still significant (partial R² = 0.06, *F*_1,84_ = 5.5, *p* = 0.021). However, most of the butterflies were sampled below 1500 m, and only individual cold adapted Pierinae from higher elevations (*Catasticta* at 2390 m and *Tatochila* at 3350 m) (Supplementary Figure S2A).

In contrast, butterflies from Germany were resampled from the same locations at different times of the year between March and September. Time of sampling (calendar week) showed a quadratic relationship with temperature (adj. R^2^ = 0.82) but was a better predictor of Shannon diversity than temperature in the same linear model (Supplementary Figure S2B). When excluding *Aglais* as low biomass outgroup the full model explained 33% of the variation in Shannon diversity (*F*_6,62_ = 6.5, *p* < 10^-4^), which could be mainly accounted to calendar week (partial R² = 0.19, *F*_1,61_ = 14.9, *p* = 0.0003), while host subfamily (partial R² = 0.04, *F*_4,62_ = 0.6, *p* = 0.63) nor temperature (partial R² = 0.02, *F*_1,62_ = 1.5, *p* = 0.23) contributed significantly. The outcome was similar when tested with host genus instead of host subfamily. This indicates that diversity of the adult butterfly microbiota shows a seasonal increase over the course of the year, independent from temperature or host taxon turnover.

For the complete dataset from both countries, we used temperature as a universal environmental predictor. Although microbial load showed an apparent quadratic relationship with temperature (Supplementary Figure S2C), this effect was not significant after accounting for host subfamily (partial R² = 0.02, *F*_2,158_ = 1.9, *p* = 0.16), whereas host subfamily explained a significant proportion of the variation (partial R² = 0.13, *F*_6,158_ = 3.9, *p* = 0.0011). For all diversity estimates, we always accounted for host subfamily when evaluating the influence of temperature on microbial richness, Shannon diversity and inverse Simpson diversity. A full model including host subfamily and temperature explained 36% of the variation in Shannon diversity (adj. R^2^ = 0.36, *F*_7,151_ = 13.89, *p* < 10^-13^). The main effect could be attributed to host subfamily (partial R² = 0.22, *F*_6,151_ = 7.2, *p* < 10^-6^), whereas temperature was not significant (partial R² = 0.02, *F*_1,151_ = 3.2, *p* = 0.076). Similar patterns were observed for microbial richness and inverse Simpson diversity (Supplementary Figure S2D–F), for which temperature was significantly associated with diversity, but explained substantially less variation than host subfamily. Notably, distribution of host subfamilies was strongly predicted by temperature (R² = 0.48, *F*_6,152_= 25, *p* < 10^-15^). This indicates that host taxonomic identity plays a larger role in shaping adult butterfly microbiota diversity than environmental temperature per se, though the occurrence of host taxa is temperature dependent.

### 3.3 Host taxonomic identity largely explains microbial community composition

Butterfly subfamilies showed significant differences in beta dispersion (*F*_6,160_ = 7.94, *p* < 0.0001), which was even more pronounced on host tribe level (*F*_11,155_ = 15.55, *p* < 0.0001) (Figure 2A). Differences among microbial community compositions were further tested by univariate PERMANOVA based on Bray-Curtis dissimilarities. As taxonomic ranks were all nested within each other, they were tested separately. Beforehand, host genera with less than two replicates were removed from the dataset. This showed increasing explanatory power (marginal R^2^) with finer taxonomic resolution: Variance explained 3.6% by host family (*F*_1,159_ = 5.8), 18% by host subfamily (*F*_6,154_ = 5.7), 25% by host tribe (*F*_10,150_ = 4.9) (Figure 2B), 33% by host genus (*F*_17,143_ = 4.1) (Supplementary Figure S3A) and 40% by host species (*F*_28,132_ = 3.1, all *p* < 0.0001). Host genera differed significantly in beta dispersion (*F*_17,143_ = 6.4, *p* < 0.0001) (Supplementary Figure S3B), but there were no such differences observed between countries (*F*_1,159_ = 1.2, *p* = 0.28) or sampling locations (*F*_3,157_ = 1.96, *p* = 0.12). Univariate PERMANOVA on these geographic predictors showed overall lower explanatory power (marginal R^2^): 6% by country (*F*_1,159_ = 9.9) and 8% by location (*F*_3,157_ = 4.5, all *p* < 0.0001).

**Figure 2.**
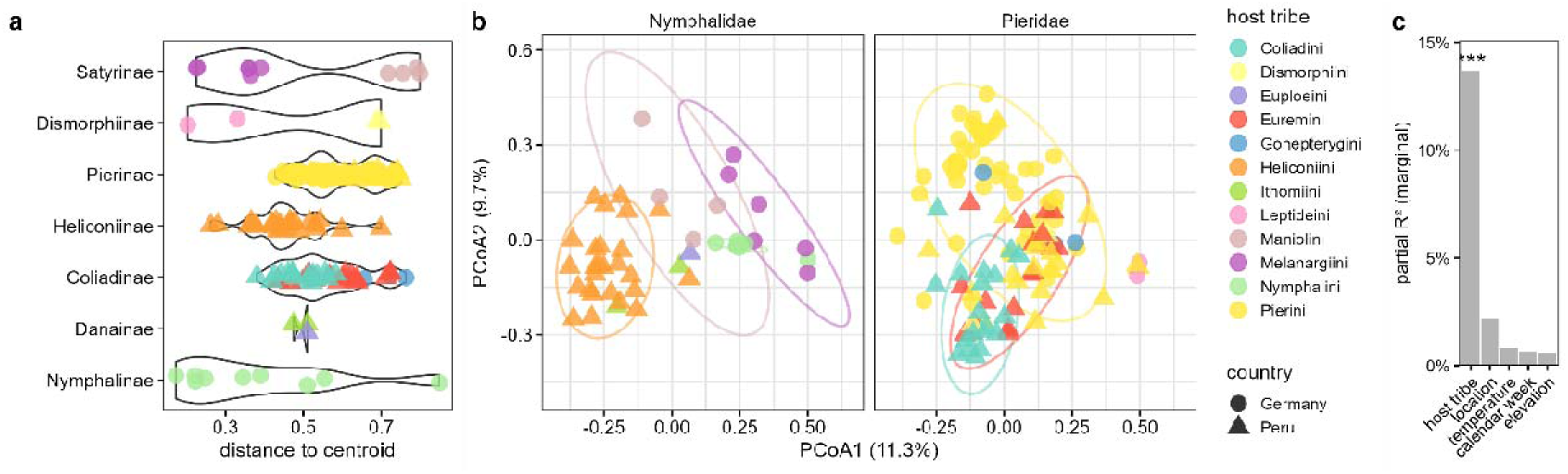
Host taxonomic identity largely explains dissimilarity of the adult butterfly microbiota composition. (**a**) Beta dispersion of the adult butterfly microbiota differs among host subfamilies (groups) and host tribes (colors). (**b**) Beta diversity of the adult butterfly microbiota using PCoA based on Bray-Curtis distances. Samples are colored by host tribe and panels faceted by host family for clarity. (**c**) Host tribe explained more variation (partial R^2^) than environmental predictors and remained the strongest predictor of microbial community composition after excluding host groups with low replication (n < 10), retaining only Satyrinae, Pierinae, Heliconiinae, and Coliadinae.

When including host taxonomy, geographic and environmental predictors in the same multivariate model we used host tribe level to avoid collinearity among geographic predictors. As there was little autocorrelation between temperature, elevation and calendar week in the full dataset (R² < 0.01), we use them together in the same model. As further robustness measure we removed host groups with low replication (n < 10) or high multivariate dispersion (i.e. Dismorphiinae, Danainae and Nymphalinae). The full model with only Satyrinae, Pierinae, Heliconiinae, and Coliadinae and explained 26% of the variation in microbial community composition (*F*_12,134_ = 3.9, *p* < 0.001), while the majority was accounted to host tribe (marginal R^2^ 0.14, *F*_6,134_ = 4.1, *p* < 0.001) (Figure 2B,C), all other environmental factors did not contribute significantly to community composition (location: marginal R^2^ 0.02, *F*_3,134_ = 1.3, *p* = 0.069; elevation: marginal R^2^ 0.006, *F*_1,134_ = 1.1, *p* = 0.35; calendar week: marginal R^2^ 0.007, *F*_1,134_ = 1.2, *p* = 0.25; temperature: marginal R^2^ 0.008, *F*_1,134_ = 1.5, *p* = 0.09) (Figure 2C), indicating that the predominance of host taxonomy was not driven by sparsely replicated groups.

In addition, we performed variance partitioning to separate the unique contribution of host taxonomic identity and environmental or geographic predictors, as well as their shared fraction. For this we used again the reduced dataset containing only Satyrinae, Pierinae, Heliconiinae, and Coliadinae, and combined temperature, calendar week, elevation and location into a single environmental predictor. This confirmed the stronger contribution of host identity as a predictor of microbial community composition compared to environment influences. Host tribe showed a unique contribution (adj. R²) of 10.3%, with a shared fraction of 5.3% and a unique contribution of the combined environmental predictors of only 4.8%. The effect in variance partitioning becomes more pronounced when using only temperature as environmental predictor (adj. R² of host tribe 12.8%, shared fraction 2.8%, unique temperature contribution 0.7%). Therefore, host taxonomic identity remained the strongest predictor in explaining the observed variation in microbial community composition than environmental and geographic factors combined.

### 3.4 General structure and composition of the adult butterfly microbiota

The microbial community composition of adult butterflies showed a high abundance and prevalence for lactic acid bacteria (LAB, order Lactobacillales rel. ab. 21.6%), enteric bacteria (order Enterobacterales rel. ab. 21.7%) and acetic acid bacteria (AAB, order Rhodospirillales rel. ab. 16.3%) (Figure 3A). Besides these, the adult butterfly microbiota contained host associated groups like Orbales, Rhizobiales & Rickettsiales (cumulative rel. ab. of 18.0%). The eight top prevalent bacterial orders made 85.6% rel. abundance of the bacterial community across all samples (Figure 3B).

**Figure 3.**
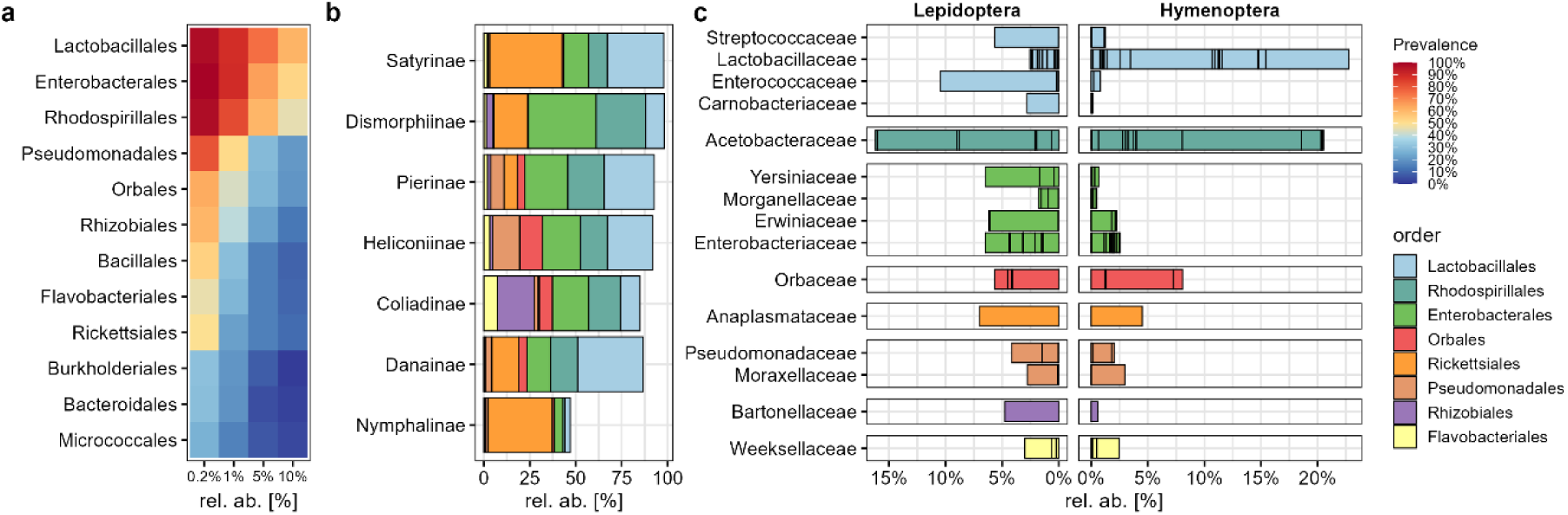
Compositional similarity of the adult butterfly microbiota across butterfly subfamilies and in comparison to bees. (**a**) Heatmap of most prevalent and abundant bacterial orders among all butterfly samples. (**b**) Microbial community composition grouped by butterfly subfamily showing the eight major bacterial orders which account for 85.6% relative abundances across all samples. (c) Relative abundances of major bacterial families from the microbiota of Lepidoptera compared to sympatric Hymenoptera from Peru (Apinae and Halictinae). Only top 16 families from butterflies are shown. Bars are colored by bacterial order and subdivided into bacterial genera.

Notably, several of the bacterial orders contained only a single bacterial family which was in turn often dominated by a single genus (Figure 3C). This was the case for insect host associated groups like Weeksellaceae (*Apibacter*), Bartonellaceae (*Bartonella*), Anaplasmataceae (*Wolbachia*) and Orbaceae (*Orbus*). Similar, Acetobacteraceae were dominated only by two genera (*Asaia* & *Commensalibacter*). Also, within the Lactobacillales most families were dominated by a single genus i.e. Enterococcaceae (*Enterococcus*), Streptococcaceae (*Lactococcus*) or Carnobacteriaceae (*Carnobacterium*). For enteric bacteria this pattern was less obvious, but some families contained also abundant genera i.e. Erwiniaceae (*Pantoea*) or Yersiniaceae (*Rahnella*), while others were more diverse.

By contrast, the family Lactobacillaceae showed only minor abundance in the Lepidoptera microbiota, with no dominant genus, but were the most abundant microbial family in the Hymenoptera dataset, comprising mainly *Apilactobacillus*, *Lactobacillus* and *Fructobacillus* (Figure 3C). Hymenoptera differed also in Acetobacteraceae, dominated by *Bombella* alongside *Asaia* and *Commensalibacter*, and in Orbaceae with the main genus *Gilliamella*. Although adult butterflies shared several dominant microbial groups with wild bees, they showed varying proportions and substantial differences in the enriched bacterial genera (Figure 3C).

### 3.5 The common genus core microbiota of adult butterflies

To identify the most abundant and prevalent bacterial genera, we performed a threshold based core analysis on the entire dataset (minimum of 20% prevalence at 1% rel. ab.), which resulted in 12 common core genera enriched in adult butterflies (Supplementary Figure S4AB). These include the LAB *Enterococcus* (Enterococcaceae) & *Lactococcus* (Streptococcaceae) showing 68% and 37% prevalence, AAB *Asaia* & *Commensalibacter* (Acetobacteraceae) with 48% and 44% prevalence, enteric bacteria *Pantoea* (Erwiniaceae) & *Rahnella* (Yersiniaceae) with 41% and 28% prevalence, as well as *Bartonella* (Bartonellaceae) 33%, *Acinetobacter* (Moraxellaceae) 24%, *Orbus* (Orbaceae) 26%, *Wolbachia* (Anaplasmataceae) 20%, *Apibacter* (Weeksellaceae) 21% and *Entomomonas* (Pseudomonadaceae) with 20% prevalence, at 1% rel. abundance. (Supplementary Figure S4A–C). Notably, several of the remaining non-core genera belonged to the same bacterial groups e.g. LAB (*Carnobacterium*), enteric bacteria (*Cedecea*, *Raoultella, Serratia* and *Klebsiella*) or Orbacea (*Frischella*), but showed more host-specific occurrence in only individual butterfly genera (Supplementary Figure S4D) or certain butterfly subfamilies (Supplementary Figure S5).

To place these common core taxa into broader taxonomic context, they were shown together with the 80 most abundant genera in a phylogenetic tree. Common core genera stand out due to their high abundance and prevalence across all butterfly samples (Figure 4). For comparison, the same threshold was applied to the bee dataset, identifying 15 common core genera that showed high prevalence in the sympatric Hymenoptera (black dots in Figure 4, Supplementary Figure S6). Comparison of both core sets identified six genera that occurred only in the common core of Lepidoptera: *Orbus*, *Enterococcus*, *Rahnella*, *Entomomonas, Bartonella,* and *Lactococcus* (Supplementary Figure S7A). Six genera showed a taxonomic overlap between the common cores of Lepidoptera and Hymenoptera: *Asaia*, *Pantoea*, *Commensalibacter*, *Wolbachia*, *Apibacter,* and *Acinetobacter* (Supplementary Figure S7B). The remaining nine genera occurred only in the common core of Hymenoptera: *Pseudomonas*, *Fructobacillus*, *Snodgrassella*, *Gilliamella*, *Lactobacillus*, *Bifidobacterium*, *Apilactobacillus*, *Bombella*, and *Pectinatus* (Supplementary Figure S7C).

**Figure 4.**
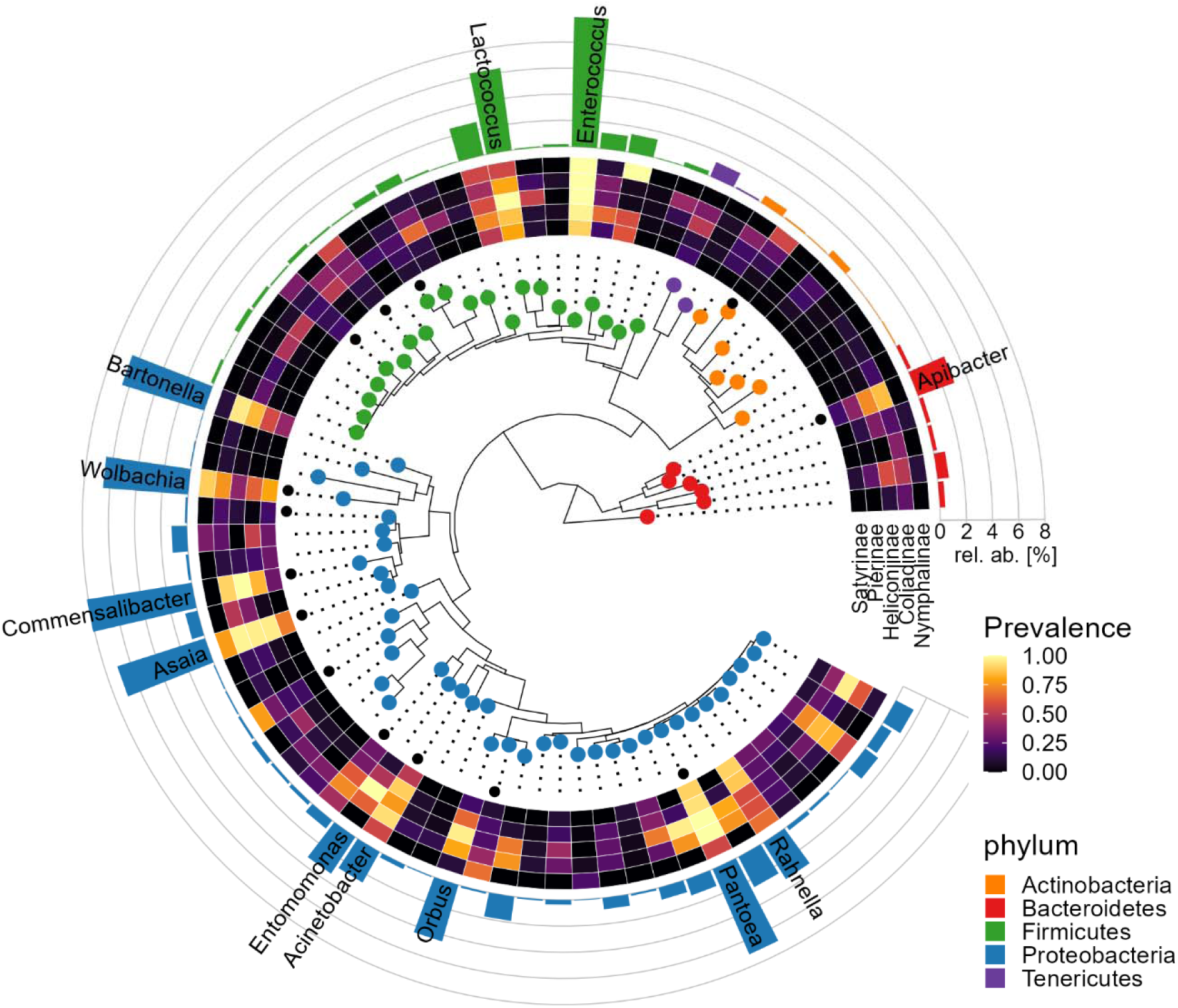
Taxonomic placement of the common core microbiota of adult butterflies. Phylogenetic grouping (SINA RAxML tree) and relative abundance of the 80 most abundant bacterial genera across all butterfly samples (represent about 98% of the total dataset). Only the common core taxa are shown with name, which show high prevalence in most butterfly subfamilies (only subfamilies with > 8 samples shown). Black dots indicate common core genera from a sympatric wild bee dataset for comparison, showing an overlap for *Apibacter*, *Wolbachia, Commensalibacter, Asaia, Acinetobacter,* and *Pantoea*. For a full genus-level annotation see Supplementary Figure S6.

### 3.6 Common core taxa distribution among countries and subfamilies

The distribution of the 12 common butterfly core genera was not equal between countries (Figure 5A). *Rahnella, Enterococcus,* and *Asaia* were both more abundant in samples from Germany (adj. *p* < 0.05), whereas *Wolbachia*, *Lactococcus*, and *Acinetobacter* were equally found in samples from both countries (adj. *p* > 0.05). On the other hand, *Orbus*, *Commensalibacter*, *Apibacter*, *Pantoea*, *Entomomonas*, and *Bartonella* were more abundant in samples from Peru (adj. *p* < 0.001) (Supplementary Figure S8). For some common core taxa, the same ASVs appear to be shared between samples from both countries (i.e. *Apibacter, Enterococcus*, *Lactococcus Orbus Pantoea* & *Rahnella*) (Supplementary Figure S9). Other common core taxa exhibited a mixture of country-specific and shared ASVs (i.e. *Asaia*, *Bartonella* & *Entomomonas*), with *Acinetobacter* & *Commensalibacter* showing the highest overall ASV diversity. Only *Wolbachia* showed clearly distinct ASVs in samples from both countries (Supplementary Figure S9). Among the 24 butterfly genera within this dataset *Wolbachia* was the most common endosymbiont and dominant in five different butterfly genera from five different subfamilies (Supplementary Figure S10). Others, like *Spiroplasma,* showed overall a minor abundance and more a loose association with individual samples from Peru but were absent in samples from Germany (Supplementary Figure S10).

**Figure 5.**
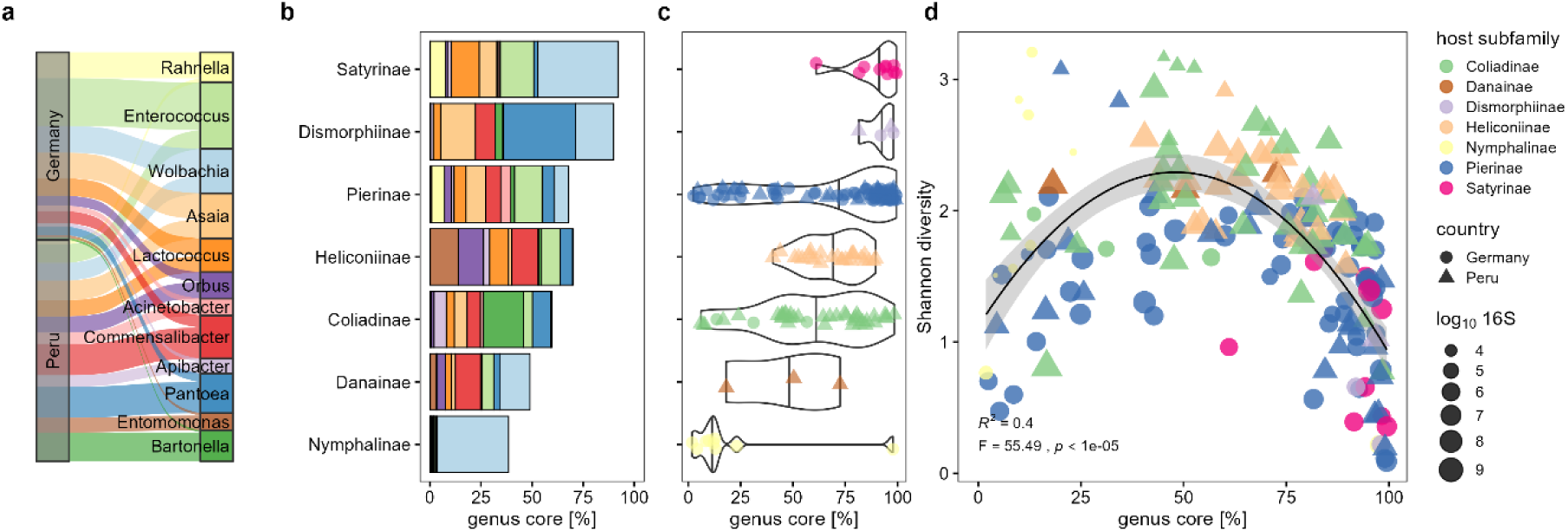
Abundance and distribution of the adult butterfly common genus core microbiota. (**a**) Country-specific distribution of the 12 common core genera. Samples from Germany (n = 76) had higher relative abundances of *Rahnella, Enterococcus,* and *Asaia,* while samples from Peru (n = 91) contained higher abundance of *Orbus Commensalibacter, Apibacter, Pantoea, Entomomonas & Bartonella*. (**b**) Mean relative abundance of the 12 common core genera among butterfly subfamilies (average abundance of 67% across subfamilies). Color legend shared with panel A. (**c**) Distribution of the common genus core among individual butterfly samples within each subfamily (see Supplementary Figure S11 for details). (**d**) Genus core abundance and microbial load showed a quadratic relationship with Shannon diversity.

Subsetting the dataset to the 12 common core genera highlights their contribution to the total relative abundance across the different butterfly subfamilies (Figure 5B). The cumulative amount of common core genera ranged from 92% for Satyrinae, 90% for Dismorphiinae, 68% for Pierinae, 70% for Heliconiinae, 60% for Coliadinae, 49% for Danainae to 38% for Nymphalinae (Figure 5B). However, abundance distribution varied among individual samples within each host subfamily (Figure 5C). There was a high variation in core genus abundance among conspecific individuals, ranging from nearly 0% to 100% as seen for *Pieris* (Supplementary Figure S5A, Supplementary Figure S11A). The low biomass outgroup *Aglais* (Nymphalinae) showed overall the lowest common core abundance, while *Polygonia* from the same subfamily showed even the highest, due to a high relative abundance of *Wolbachia* (Supplementary Figure S4C, Supplementary Figure S11A). The relative abundance of all common core genera varied significantly among butterfly subfamilies (all genera KW adj. *p* < 0.007) (Supplementary Figure S11B). Some groups like *Bartonella* occurred mainly in Coliadinae, while *Entomomonas* seemed characteristic for Heliconiinae (Figure 5B) (Supplementary Figure S11B), which explains partly the country-specific distribution.

Interestingly, genus core abundance as well as microbial load both showed highly significant quadratic relationships with Shannon diversity (overall adj. R^2^ = 0.50, *F*_4,162_ = 42, *p* < 10^-15^). Both retained independent nonlinear associations when included in the same model, with genus core abundance explaining more variation (partial R^2^ = 0.39, *F*_2,162_ = 52, *p* < 10^-15^) than microbial load (partial R^2^ = 0.18, *F*_2,162_ = 18, *p* < 10^-7^) (Figure 5D). This result provides an unexpected link between total sample diversity, total microbial abundance, and abundance of common genus core taxa. Nonetheless, this pattern could provide a basis for the mechanistic understanding of the principles during community assembly in solitary pollinators.

## 4 Discussion

Our systematic assessment of the microbiota composition from adult butterfly populations across different continents revealed that adult Lepidoptera can assemble abundant gut microbiota distinct from those of Hymenoptera. The microbiota of adult butterflies showed conserved common core taxa shared across a variety of different butterfly subfamilies, with only partial overlap with those associated with sympatric bees. Our analysis showed that the variation of microbial communities was best explained by host taxonomic placement with only little environmental influence. Previous assumptions of uniform microbial community compositions within these mostly generalist nectar feeders do not hold true for adult butterflies^44^.

### 4.1 Host taxonomic identity as best predictor of the adult butterfly microbiota

Our finding is in line with the observations by Ravenscraft, *et al.*^32^, who found a stronger influence of host species identity compared to host feeding preferences in Neotropical butterflies (with > 80% of samples from Nymphalidae including many fruit-feeding species). Similar, Hammer *et al*.^33^ described phylogenetically structured microbiomes within Neotropical Heliconiinae, but found only a minor influence of pollen-feeding. Extending on these two previous studies, our dataset included Neotropical as well as Palearctic species and a broader taxonomic range with 70% of samples within the family of Pieridae (subfamilies: Pierinae, Coliadinae & Dismorphiinae). This study represents the first cross-continental comparison of adult butterfly microbiota, and some taxonomic groups might still be underrepresented. Ongoing sampling efforts of local butterfly populations, as done within the framework of the LepEU consortium^45^, will enable a targeted pan European comparison of selected host species. Still, we found a strong influence of host taxonomic placement, while our alternative environmental predictors based on biogeographic pattern, elevation or temperature showed only minor influence on the adult butterfly microbiota. Temperature is usually an important predictor of environmental microbial diversity^46^, as well as insect abundance at the study site in Peru^47,48^. Therefore, temperature appears to influence microbiota diversity of butterflies only indirectly, by affecting the presence of host taxa. Individual cold-adapted species sampled from high elevations (*Catasticta* and *Tatochila*) or early in season (*Aglais*) showed lower microbial richness and lower occurrence of common core taxa, with predominant colonization by generalist bacteria (*Pseudomonas* or *Bacillus*) (Supplementary Figure S4D). Microbial associations of insects appear to be less specific under colder conditions, fitting to similar observations made with mosquitoes from Greenland^49^, or bees and butterflies from montane environments^42,50^.

In contrast, environmental factors and geographic locations can be dominant drivers of microbiota composition in other model insect systems. A cross-continental survey of *Drosophila* populations showed that abiotic factors explain most of the variation in microbiome diversity^51^. *Drosophila* exhibit largely neutral microbiome assembly^52,53^ with limited influence of host phylogeny^54^. However, their ecology differs fundamentally as fruit flies develop and proliferate on decaying fruits, which can be a reliable microbial pool when adults emerge. In contrast, butterflies undergo pronounced ontogenetic niche shift and nectar-feeding adults encounter a transient and variable distribution of nectar microbes^55^. Nevertheless, even fruit-feeding and nectar-feeding species showed similar microbial richness and only minor differences in microbiota composition^32^. This contrasts findings in lepidopteran larvae, which contain transient, environmental strains which often resemble that of their local food plants^16^. Although adult butterflies in this study have been collected from different climate zones, they were colonized by a consistent set of common core taxa, typically found in insect gut environments. Remarkably, we found that several core taxa ASVs were identical in samples from both countries. This suggests that host-filtering is a dominant factor driving butterfly microbiota, resulting in the consistent enrichment of specific microbial groups from diverse environmental sources. At the same time, the widespread occurrence of these taxa indicates that they possess general traits that facilitate persistence and successful colonization of various insect gut environments.

### 4.2 Insect-associated bacteria as adult butterfly common core taxa

We identified twelve common core genera, all of which were also reported as prevalent groups in all previous studies of Neotropical butterflies^28,32–35^. *Wolbachia* is probably the most common endosymbiont reported in Lepidoptera^37,38^ and showed host-genus specific distribution due to its mainly vertical transmission mode. Other endosymbionts like *Spiroplasma* occurred in lower frequency in our dataset, but were also previously reported in *Heliconius*^33–35^. Interestingly, *Heliconius* harbors either *Spiroplasma* or *Kinetoplastibacterium*, but never both within the same specimen (see Supplementary Figure S10). Another key taxon is *Orbus*, which was originally described as isolate from an adult butterfly *Sasakia charonda* (Apaturinae)^39^. Later, the family of Orbaceae became mainly known as major gut symbiont of bees^56^, but has recently been obtained as well from drosophilid and tephritid fruit flies^57^. *Entomomonas* shows broad insect-associations and seems to be another characteristic core taxon of *Heliconius* butterflies^33,58^.

### 4.3 Overlap and distinction of the lepidopteran and hymenopteran microbiota

The characteristic social core microbiota of corbiculate bees (Hymenoptera: Apinae) was, as expected, largely missing in solitary butterflies (i.e. *Bifidobacterium*, *Snodgrassella*, *Gilliamella* & *Lactobacillus*) (Supplementary Figure S7C). However, some common core genera observed in adult butterflies showed an overlap with the common core of sympatric hymenopteran pollinators: *Apibacter*, *Acinetobacter, Bartonella* and *Commensalibacter* are also commonly associated with bees. *Apibacter* and *Acinetobacter* are considered as environmentally acquired strains, but widely found in the microbiota of true social bees (*Apis*, *Bombus*)^59,60^. In our analysis, they showed significantly higher abundance in Hymenoptera than in Lepidoptera (Supplementary Figure S7B). *Bartonella* (Alpha-1) and *Commensalibacter* (Alpha-2.1) occur in lower amounts in honey bees^61^, but can become abundant during overwintering^62^. Likewise, *Commensalibacter papalotli* has been isolated from overwintering monarch *Danaus plexippus* (Nymphalidae) from Mexico^40^. Other *Commensalibacter* isolates have been obtained from different Nymphalidae and Pieridae butterflies in North America, Europe and Asia^63–65^, indicating that this common core genus occurs across a broad range of butterfly species worldwide. Other AAB, such as *Asaia,* show generally very broad insect associations^66^, whereas *Bombella* is characteristic of bees. Overall, AAB represent dominant members of the microbiota of Neotropical orchid bees (Euglossini) and stingless bees (Meliponini)^3,42^. *Pectinatus* was the only genus in our dataset unique to bees and was entirely absent from butterflies (Supplementary Figure S7C). This genus has been mainly reported from Meliponini^3,67^ and giant honey bees^68^ and is considered a strict anaerobe. Its absence from adult butterflies suggests that their gut environment may favor more aerotolerant bacteria and may be less oxygen-depleted than the bee hindgut.

Another interesting distinction between the microbiota of Hymenoptera and Lepidoptera can be seen in lactic acid bacteria (LAB). Lactobacillaceae are a prominent bacterial group of social bees (*Apis*, *Bombus, Melipona*)^67,69,70^ and comprise a high diversity of genera, including *Lactobacillus* and *Apilactobacillus*. Ground nesting bees (Colletidae & Halictidae) can even be dominated by a single strain of *Apilactobacillus*^71,72^. In contrast, these specialized fructophilic and acid-tolerant LAB are underrepresented and nearly absent from butterflies. Though it might be intriguing to hypothesize that Lactobacillaceae are acquired via pollen feeding, they were not particularly enriched in *Heliconius*, which represents the only pollen-feeding butterfly group in our dataset. Only *Tithorea* (Danainae) harbored considerable abundances of *Lactiplantibacillus*, *Secundilactobacillus*, and *Levilactobacillus*.

Instead, most adult butterflies show high abundance and prevalence for *Enterococcus* (Enterococcaceae) and *Lactococcus* (Streptococcaceae), which represent LAB with broader or more moderate pH tolerance, respectively. *Enterococcus* is a physiologically highly versatile bacterium and associated with a broad variety of different hosts giving them a character as generalist^73^. They are also frequently reported as dominant member of the larval gut microbiota of certain moth families, providing potential antimicrobial and antiparasitic functions^31^. This makes *Enterococcus* a signature taxon with a prominent role in lepidopteran life cycle. *Enterococcus* is usually rare in Hymenoptera, while *Lactococcus* can occur in some bees. In stingless bees (*Melipona* spp.) a related, flower-associated genus *Floricoccus* (Streptococcaceae) has been described recently^67,69^.

### 4.4 Environmental acquisition and microbial community assembly in butterflies

Adult butterflies are exposed to a highly stochastic microbial pool of nectar microbes, driven by passive dispersal and random visitation of pollinators^55^. Nevertheless, insects can acquire host-specific microbiota from ephemeral environmental sources. Ground-nesting sweat bees (Halictidae) can selectively enrich a single *Apilactobacillus* strain from pollen provision, but will lose them under artificial rearing conditions^72^. Even obligate symbioses can evolve from environmental acquisition, which may make these associations more vulnerable to disturbances, while providing ecological flexibility that can foster host adaptations to changing environmental conditions^74,75^. Unlike social insects, butterflies are exposed to environmental microbes immediately after emergence and lack persistent social interactions and direct transmission routes such as trophallaxis, resembling a lifestyle more comparable to that of solitary bees^5^. Nevertheless, aggregations at shared feeding sites may increase the probability of microbial exchange among conspecifics^76,77^. Many butterfly species aggregate when performing mud puddling behavior to obtain additional salts and amino acids from moist soil, dung or even carcasses^78^. This has been frequently observed in the tropics but also in temperate regions among various Pieridae and Nymphalidae, and is an intriguing but yet untested potential microbial source for adult butterflies^32^.

Nonetheless, the composition of the adult butterfly microbiota largely resembles that of insects feeding on sugar-rich substrates^66^, and contains taxa associated with floral nectar, such as *Acinetobacter*, also common in other pollinators^60^. We measured for most butterflies microbial loads around 10^6^ to 10^8^ 16S copies which were in the range of previous reports from Heliconiini^33^ or stingless bees^3^, while others have reported even higher amounts from butterflies^32^. Notably, similar microbial quantities have been reported during microbiome assembly in honey bees^79^ The gut environment of nectar-feeding butterflies is probably highly selective, enriching microbes adapted to high sugar concentrations, while host-filtering others. Though, it cannot be excluded that other ecological or physiological factors may contribute to this (e.g. host immune parameter)^4^. The observed relationship between Shannon diversity and common genus core abundance (Figure 5D) seems, at first, surprising but could provide insights about the microbiota assembly process. Initial colonization may be driven by stochastic acquisition and priority effects^4^, resulting in communities dominated by non-core bacteria. Secondary microbial uptake may initially increase both microbial diversity and the representation of core taxa. With a transition toward a more established and core-dominated community, diversity may subsequently decline due to the increasing dominance of core taxa. Thus, core microbiota dominance may provide an indicator of microbiome establishment that could complement microbial load measurements. Future studies with more extensive replication of individual butterfly species under comparable environmental settings will be required to disentangle the relative contributions of deterministic and stochastic processes during butterfly microbiome assembly. In addition, we identified a seasonal component, indicating that microbial diversity changes over the course of a year independent from temperature or host turnover. Considering ongoing climate-driven range shifts documented for butterflies globally^80^, consequences for host-microbiome associations have yet to be determined.

The low bacterial biomass and almost lack of a common core microbiota in the early spring sampled *Aglais* butterflies is not considered an artifact, but rather evidence that these butterflies have not yet acquired a host-specific microbiota and start ‘empty’ during their very first flights in March. This is consistent with the physiology of overwintering adults, as they usually purge their entire gut content before diapausing to avoid the presence of ice nucleators during hibernation^81^. Spore forming Bacilli are likely only inactive and transient gut members that can become dominant reads in metabarcoding studies when the amount of active bacteria is low^82^. But even *Aglais* can assemble common core taxa as revealed in another study that sampled later in season (July – October)^83^. Still, our data suggests that overwintering adults do not retain gut microbes from the previous year but rather acquire them newly when flying in spring.

### 4.5 Functional implications of the adult butterfly-associated microbiota

Even nectar-feeding adult butterflies need amino acid supplementation in their diet^84^, which could be facilitated by symbiotic bacteria^85^. Several enteric bacteria commonly found in insect guts have potential nitrogen fixing abilities, including the butterfly core members *Rahnella* & *Pantoea*^86^. However, direct functional evaluations of the adult butterfly microbiota are rare. Antibiotic treatment shortened adult life span in the Indian Cabbage white *Pieris canidia* (Pieridae)^87^, and a similar correlation between bacterial colonization and adult longevity was observed in *Speyeria mormonia* (Nymphalidae)^63^. In *Pieris brassicae* (Pieridae), experimental depletion of the adult microbiota revealed an unexpected trans-generational phenotype on the offspring, influencing larval biomass gain and immune responses during a host plant shift^29^. Maternal and paternal effects on offspring performance are well documented across butterfly species^88^, yet the potential role of adult microbial load in these phenomena has not been explicitly considered.

### 4.6 Conclusion and Outlook

Our findings provide insights into how host taxonomic identity and environmental factors shape the adult butterfly microbiota. It becomes obvious that even basic principles about the assembly and functional relevance of the adult butterfly microbiota remain poorly investigated aspects of Lepidopteran ecology. More extensive samplings across a broader taxonomic range among local populations could help to consolidate these findings and further disentangle environmental influences from host-specific effects. Our work provides a foundation to test how butterfly age or season influences core taxa assembly and diversity over time. Such work could help to evaluate the extent to which anthropogenic influences could potentially disturb adult butterfly microbiota in natural context.

## 5 Material and Methods

### 5.1 Sample collection and environmental temperature

Samples from Peru were collected between November 2022 and May 2023 within the Kosñipata valley from two main locations (provinces Paucartambo and Manu) as part of the ANDIV project (https://www.andiv.biozentrum.uni-wuerzburg.de/). Sampling sites include nine sublocations located along an elevational gradient reaching from 245 m up to 3350 m above sea level^89^. Target groups were ‘whites’ and ‘sulfurs’ from the Pieridae family as well as *Heliconius* (Nymphalidae) as reference group. Butterflies were sampled by hand using a sweep net and directly dissected at local research stations. Entire abdomens were stored in DNA/RNA Shield buffer (Zymo Research) or in 100% Ethanol for preservation and transport at ambient temperatures. From each specimen a single leg was stored in 100% Ethanol for DNA barcoding and species identification at the Canadian Centre for DNA Barcoding (CCDB) in Guelph (Canada). The obtained barcodes of the cytochrome oxidase (COI) sequences are accessible on https://portal.boldsystems.org/ (see Data availability section). Wings and remaining body parts were desiccated and stored in glassine envelopes to aid with species identification. All butterflies were classified up to genus level, and if possible, to species level. For comparison we included a sympatric wild bee microbiota dataset from Peru^42^, which was processed under the same laboratory conditions. European butterfly species were collected near Nuremberg in middle Franconia in Germany. The two major sampling locations are approx. 26 km apart at 310 m and 350 m above sea level, respectively, and include four sublocations. Butterflies were collected between March and September in three consecutive years (2021,2022 and 2023) by hand using a sweep net and identified to species level morphologically. Most samples were from two common species *Pieris napi* and *P. rapae* (Pierinae). Samples were stored at -20°C and abdomen dissected under laboratory conditions. Storage conditions had no influence on microbial community composition when accounting for host tribe and location within the same model (PERMANOVA Storage: partial R^2^ = 0.006, *F*_1,146_ = 1.17, *p* = 0.27). A total of 199 butterfly samples were analyzed, 104 from Peru and 95 from Germany.

The average air temperature in the seven days before sample collection was taken as the sampling timepoint ‘temperature’ for each individual sample. The environmental temperatures at each sampling plot in Peru were directly measured with temperature loggers, showing a temperature gradient from 8.1°C to 26.1°C across the elevational gradient^48^. For locations within Germany, measured temperatures for each sampling timepoint were taken from https://meteostat.net, showing a temperature gradient from 6.7°C to 24.8°C across season.

### 5.2 DNA Isolation, PCR conditions and library preparation

DNA isolation of entire abdomen was performed using ZymoBIOMICS™ 96 kits (Zymo Research). Bead beating of the ZR BashingBead™ lysis tubes was performed at 2500 rpm for 30 min on a multi-tube vortex mixer (Ohaus) and an additional Lysozyme digestion step with 50 U/µl Ready-Lyse Lysozym solution (Biozym) for 15 min at 25°C included. The ZymoBIOMICS Microbial Community Standard as well as ZymoBIOMICS Spike-in Control I (Zymo Research) was used as positive control. On each extraction plate a minimum of six negative extraction controls (NECs) were included. DNA isolation was performed with an Integra Assist Plus pipetting robot placed under a vertical laminar air flow cabinet AirClean 600 (Syngene). DNA was eluted in a volume of 50 µl and stored at -20°C until library preparation.

The PCR for metabarcoding was performed according to the dual-indexing strategy by Kozich *et al*.^90^ and the pipetting scheme from Sickel *et al*.^91^. Amplification target was the V4 region of the 16S SSU rRNA gene. The primers contained Illumina adaptor P5 (forward) or P7 (reverse), a unique 8 base barcode, a Pad sequence followed by the earth microbiome primer sequences 515F (GTGCCAGCMGCCGCGGTAA) and 806R (GGACTACHVGGGTWTCTAAT)^92^. PCR reactions were performed in triplicates, with 1 µl of DNA template in 10 µl final reaction volume using Thermo Phusion Plus PCR-Mastermix (Thermo Fisher Scientific) and 0.33 μM of each primer. The PCR protocol consisted of an initial activation at 98°C for 30 sec, followed by 30 cycles of 98°C for 10 sec, 55°C for 10 sec, 72°C for 30 sec and a final extension at 72°C for 5 min. PCR amplicons were assessed on precast E-Gel™ 96 Agarose Gels with SYBR Safe on a E-Gel Power Snap Plus Electrophoresis System (Thermo Fisher Scientific). Samples with no visible band on the agarose gel were subjected to a second PCR. Samples for which repeated amplification remained negative were later excluded during analysis (n = 17 samples). Exceptions were made for *Aglais* samples that were kept on purpose as an outgroup within the dataset (see explanation below). Amplicons were normalized using SequalPrep™ Normalization Plate Kit (Thermo Fisher Scientific) and the plate pools were concentrated using AMPure XP Beads (Beckman Coulter). A final linearization PCR was performed with each plate pool library including primer based on the Illumina adapter sequences. Each 25 µl reaction contained 11 µl of plate pool library, 0.3 µM forward primer P5 (AATGATACGGCGACCACCGA), 0.3 µM reverse primer P7 (CAAGCAGAAGACGGCATACGA) and 12.5 µl PCR mix Thermo Phusion Plus PCR-Mastermix. The cycling conditions were as follows: 98°C for 3 min, followed by one cycle of 98°C for 1 min, 50°C for 1 min and 72°C for 5 min. Final products were cleaned by AMPure XP Beads and quantified using 1×dsDNA High Sensitivity Assay on a Qubit 4 Fluorometer (Thermo Fisher Scientific). Fragment size and library quality was checked on a 2100 Bioanalyzer using High Sensitivity DNA Chip (Agilent Technologies). The final library was pooled equimolarly at 2 nM for paired-end sequencing and 5% PhiX was added to increase read quality. Sequencing was performed at the Genomics Service Unit of the LMU Biocenter on an Illumina MiSeq platform using Illumina Reagent Kit v2 (2 × 250bp).

### 5.3 Absolute quantification of 16S rRNA gene copy numbers by qPCR

Quantification of 16S rRNA gene copy numbers was performed using adapter-free versions of the 515F/806R primers. PCR reactions were performed in triplicates, with 1 µl DNA template per 10 µl final reaction on a CFX96 Touch Real-Time PCR Detection System (BioRad) using the Power Track™ SYBR Green Master Mix (Applied Biosystems). Cycling conditions followed a two-step protocol with initial activation at 95°C for 2 min, followed by 40 cycles of 95°C for 10 sec and a combined Annealing/Extension step at 60°C for 30 sec, including a final dissociation curve. Standard curves were generated from serial dilutions of the Femto Bacterial DNA Quantification Kit Standards (Zymo Research) ranging from 10 ng to 10 fg genomic DNA per reaction, corresponding to log_10_ 16S rRNA gene copy numbers of 7.15 to 1.15. Standard curves showed coefficients of determination (R^2^) > 0.99 and a mean amplification efficiency (E) of 80%. Absolute copy numbers per specimen were calculated considering a 2.5-fold dilution during DNA isolations and the final 50 µl elution volume. Non-template controls (NTCs) were used to determine the limit of detection (LOD), corresponding to a mean of 1.11 log_10_ 16S copies per reaction (equivalent to 3.21 log_10_ 16S copies per specimen).

### 5.4 Bioinformatics and metabarcoding sequence processing

Sequencing reads were processed using the custom metabarcoding pipeline available at https://github.com/chiras/metabarcoding_pipeline^93^. Paired end forward and reverse reads were joined using VSEARCH V2.21.1^94^ with a minimum overlap of 20 bp and a maximum of 10 mismatches. Reads were removed if shorter than 170 bp, contained ambiguous bases (N), or had a maximum expected error greater than 1 (maxEE = 1). Sequences were dereplicated and denoised to generate amplicon sequence variants (ASVs) using UNOISE^95^, followed by singleton removal and chimera filtering^96^. ASVs were mapped for iterative classification against the RDP (v18), Greengenes (v13.5) and SILVA (v123) database using a global alignment identity threshold of 97%. Remaining reads without taxonomic allocation were hierarchically classified against the RDP (v18) database using SINTAX^97^. Individual ASV sequences were manually validated using NCBI BLASTN v2.16.0 via the web interface and by the 16S-based identification available at https://www.ezbiocloud.net/identify. Data was further processed in R version 4.4.2^98^ using a PHYLOSEQ v1.50.0 pipeline^99^, including the packages MICROVIZ v0.12.5^100^, GGPLOT2 v3.5.1^101^, DECONTAM v1.26.0^102^, VEGAN v2.6-8^103^. Non-bacterial reads, including plant-derived chloroplast and mitochondrial sequences, and reads unassigned at phylum level were removed from the dataset, accounting for an average of 0.38% of total reads. Samples were excluded when non-bacterial reads exceed both 500 reads and 2% rel. abundance. ASVs with less than 60 reads (<0.01‰ of total reads) and genera below 600 reads (<0.1‰ of total reads) were filtered from the dataset, removing 1.1% of total reads. ASVs belonging to the microbial community standard used as positive controls were removed from the entire dataset. The resulting cleared positive controls were used together with negative extraction controls (NECs) to identify potential reagent contaminations using the prevalence based method implemented in the DECONTAM package^102^, resulting in an average removal of 0.13% of reads. Samples were evaluated for sequencing completeness rather than sequencing depth using the INEXT package v3.0.2^104^, retaining only samples with high sample coverage (SC > 98%) for analysis

The final dataset contained 167 samples with 5,792,721 total reads and a median of 34,989 reads per sample, including 951 ASVs representing 146 bacterial genera. Final read depth and sample coverage per sample with all processing steps are provided in Supplemental Table S1. Differences in sequencing depth were accounted for by total sum scaling (TSS) to obtain relative read abundances (RRA) per sample. For most analyses ASVs were collated on genus level.

*Aglais* samples showed overall the lowest sampling throughput (mean 5,245 reads) and highest proportions of filtered reads (average of 6.5% non-bacterial reads, 9.7% low abundant taxa and 3.1% removed by the DECONTAM package). Their microbiomes were mainly dominated by *Bacillus*. This was indicative for a low microbial biomass, which was confirmed by qPCR. Still, *Aglais* samples were intentionally retained in the dataset for comparative purposes as they represent an outgroup sampled at very low temperatures from an exceptionally early sampling time point. Due to their low microbial biomass and consequently low sequencing depth, they were excluded from diversity estimates to avoid any depth-related bias.

The combined Lepidoptera-Hymenoptera dataset was processed similarly. As bee samples showed substantially higher proportions of plant-associated reads, samples were excluded only when non-bacterial reads exceeded both 5,000 reads and 20% rel. abundance. The final dataset contained 872 bee samples from the subfamilies Apinae (Apini n = 69, Bombini n = 45, Euglossini n = 139, Meliponini n = 415) and Halictinae (n = 204), mainly from the tribe Augochlorini.

### 5.5 Core microbiota analysis and phylogenetic placement

Common core genera were defined based on abundance and prevalence thresholds using the core_members function implemented in the MICROBIOME package v1.28.0^105^. Given the heterogeneous assemblage of butterflies in our dataset (35 species from 24 genera, 12 tribes and 7 subfamilies), we used less stringent and more general definitions compared to other studies on single social species. A medium abundance threshold was chosen as an intermediate core definition (minimum relative abundance of 1% with prevalence of 20%), which resulted in 12 common core genera across all butterfly samples (Supplementary Figure S4B). Similar results were obtained using a high abundance definition (5% rel. abundance and 10% prevalence) which resulted in a subset of 11 common core genera. The low abundance definition (0.2% rel. abundance with 40% prevalence) recovered 10 of the 12 common core members. For direct comparison, we used the same operational definition (1% rel. abundance with 20% prevalence) for the Hymenoptera dataset, identifying 15 common core genera among the 872 sympatric bee samples representing 34 genera, 6 tribes, and 2 subfamilies.

To build the phylogenetic tree of the butterfly microbiota, the dataset was simplified to the 80 most abundant bacterial genera meeting a minimum prevalence of 5% and minimum abundance of 2000 reads, accounting for 98% of total reads in the dataset. The most abundant ASV of each genus was exported as representative sequence using BIOSTRINGS v 2.74.0^106^ and APE v 5.8^107^. ASV sequences were combined with a custom reference set and imported into SINA aligner v1.2.12 and aligned against neighboring query reference sequences from the SILVA database using minimum identity threshold of 0.95 (https://www.arb-silva.de/aligner/). A maximum-likelihood tree was constructed in RAxML using a GTR substitution model and Gamma likelihood. The resulting tree was re-imported into R, reference sequences were removed, and the tree was rooted to Chitinophagaceae as outgroup using GGTREEEXTRA v 1.16.0^108^.

### 5.6 Statistical analysis

As predictors for microbial alpha and beta diversity, we tested nested host taxonomic ranks individually (i.e. host ‘family’, ‘subfamily’ or ‘tribe’). To avoid model overfitting most tests were done with these higher taxonomic ranks or on a reduced dataset where unique genera have been removed. Collinearity among all predictors was assessed in each model using the variance inflation factor (VIF < 3), and each model was checked for aliased coefficients. As butterfly ‘genus’ was structurally confounded with ‘country’ (2 levels) and ‘sublocation’ (13 level), we did not include them in the same model. To account for this, we primarily used only higher taxonomic ranks together with ‘location’ (4 levels) as the main geographic predictor, which includes two major sampling locations per country (Paucartambo n = 67, Manu n = 24, Nuremberg n = 70 and Hersbrucker Alb n = 6). We used ‘temperature’ as a unified environmental predictor as well as ‘elevation’ above sea level for the Peru dataset and sampling timepoint (‘calendar week’) for the Germany dataset.

For alpha diversity estimations, all retained samples met the minimum sample coverage threshold (SC > 98%). We calculated observed richness, the Shannon index (*H*′) and inverse Simpson index, corresponding to Hill numbers of orders q = 0, 1, and 2, respectively^109^. Predictors were tested with linear models (lm) and the Anova function with Type II sums of squares from the CAR package v3.1-3^110^. All tests were two-sided. Model residuals were assessed for normality by Shapiro-Wilk tests and Q-Q plots. Adjusted R^2^ values were used to quantify overall model fit and multiple predictors were retained in the models if they improved support according to Akaike Information Criterion (AIC). The contribution of each individual predictor was estimated using Type II sums of squares, and expressed as partial R², verified by partial eta-squared values from the EFFECTSIZE package v1.0.1^111^.

Beta diversity was visualized using Principal Coordinates Analysis (PCoA) based on Bray-Curtis distances from relative abundance data. In the combined Lepidoptera-Hymenoptera dataset, non-metric multidimensional scaling (NMDS) was based on Bray-Curtis dissimilarities of square-root-transformed relative abundance data. Differences in community composition were tested using a PERMANOVA based on a Bray-Curtis distance matrix via the adonis2 function, and by permutational tests on group dispersion (distance to the centroid), both with 999 permutations as implemented in the VEGAN package ^103^. PERMANOVA was conducted using marginal (Type III-like) sums of squares to assess the contribution of each predictor (partial R²) after accounting for all other terms in the model. To evaluate the unique and shared fractions of explained variation, we performed variance partitioning on a Hellinger transformed community matrix using the varpart function from the VEGAN package. This provides the unique fraction explained by host factors versus environmental factors (adjusted R²) as well as the variation shared among predictors. As a robustness analysis we used a reduced dataset retaining only well replicated subfamilies (n ≥ 10). Differences in abundance distribution of common core taxa among host groups or countries was assessed using individual Wilcoxon rank-sum Tests or Kruskal-Wallis-Tests with Benjamini–Hochberg correction accounting for multiple comparisons.

## 6 Data availability

The raw sequencing reads generated in this study have been deposited in the NCBI Sequence Read Archive (SRA) under BioProject PRJNA1426605 (https://www.ncbi.nlm.nih.gov/bioproject/1426605). The obtained barcodes of butterfly species are accessible via the Bold Systems portal (https://portal.boldsystems.org/) via the process IDs: ANDIV1591-23 to ANDIV1615-23, ANDIV1901-23 to ANDIV1928-23 and ANDIV11933-24 to ANDIV11983-24. Bee microbiome data from Pinos *et al*^42^ is available under BioProject: PRJNA1294982.

## 7 Code availability

The full analysis pipeline, including all code and data necessary to reproduce the 16S sample processing, figures and statistical analyses, is available on GitHub (https://github.com/micromole/butterfly-core-microbiota).

## 8 Acknowledgement

The authors would like to thank Uschi Schkölziger and Anika Schopf for technical help during DNA isolation, library preparation and qPCR as well as Andreas Brachmann for sequencing. We also thank Constantin Stefan for assistance with field sampling in Peru. We are grateful to the National Service of Natural Protected Areas (SERNANP) and the National Forestry and Wildlife Service (SERFOR) of Peru for permissions for field work. Samples have been collected under the SERFOR permit number AUT-IFS-2022-102 (N° D000178-2022-MIDAGRI-SERFOR-DGGSPFFS-DGSPFS) as well as by SERNAP permit N°011-2022-SERNANP-JEF. Export was done in accordance with permit N° 003756-SERFOR.

## 10 Author contributions

Conceptualization, A.W.; Methodology, A.W., A.K.; Investigation, A.W., A.P., Y.C.-C., K.L.H., P.A.-A., F.Y.; Sample acquisition, A.W., A.P., Y.C.-C.; Data curation, A.W., K.L.H.; Data analysis, A.W.; Writing – Original Draft, A.W.; Writing –Review & Editing, A.W., A.P., Y.C.-C., K.L.H., P.A.-A., F.Y., I.S.-D., M.K.P., G.B., A.K.; Funding Acquisition, A.W., I.S.-D., M.K.P., G.B., A.K.; Resources, A.K., F.Y.

## 12 Supplementary Material

**Supplementary Figure S1.**
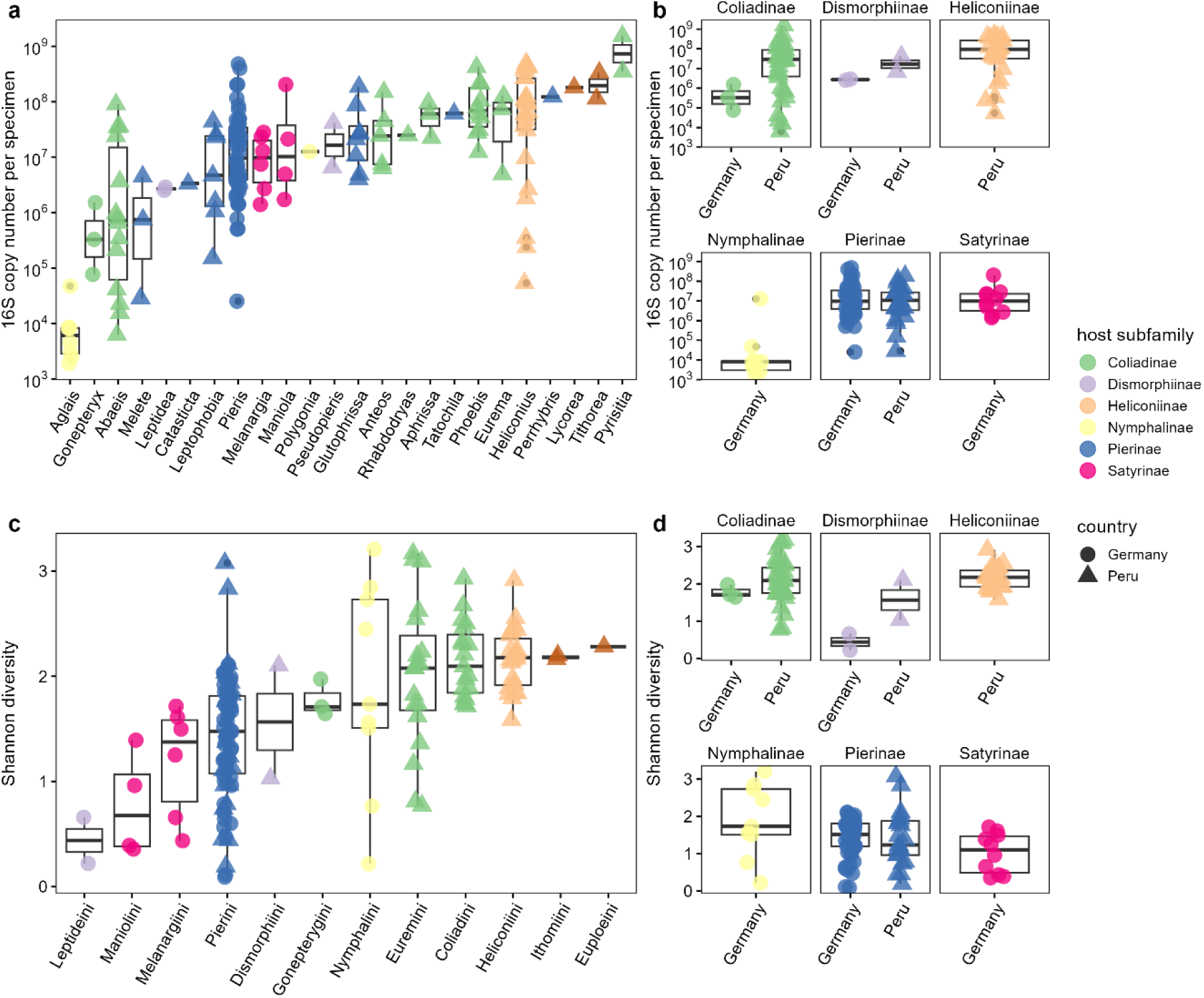
Bacterial load and diversity across butterfly host taxa and countries. (**a**) Total bacterial load estimated by 16S rRNA gene copy number qPCR per butterfly genus. (**b**) Bacterial load did not differ between countries in individual subfamilies. Wilcoxon rank-sum tests: Coliadinae (*p* = 0.051), Dismorphiinae (*p* = 0.33) and Pierinae (*p* = 0.81). (**c**) Shannon Index grouped by host tribe and colored by host subfamily. The full model explained 30% of variation in Shannon diversity, with the main effect accounted to host tribe (partial R² = 0.24, *F*_11,154_ = 4.3, *p* < 10^-4^), there was no effect of country (partial R² < 0.01, *F*_1,154_ = 0.03, *p* = 0.87). (**d**) Shannon Index did not differ by country in individual Wilcoxon rank-sum tests: Coliadinae (*p* = 0.12), Dismorphiinae (*p* = 0.33) and Pierinae (*p* = 0.62).

**Supplementary Figure S2.**
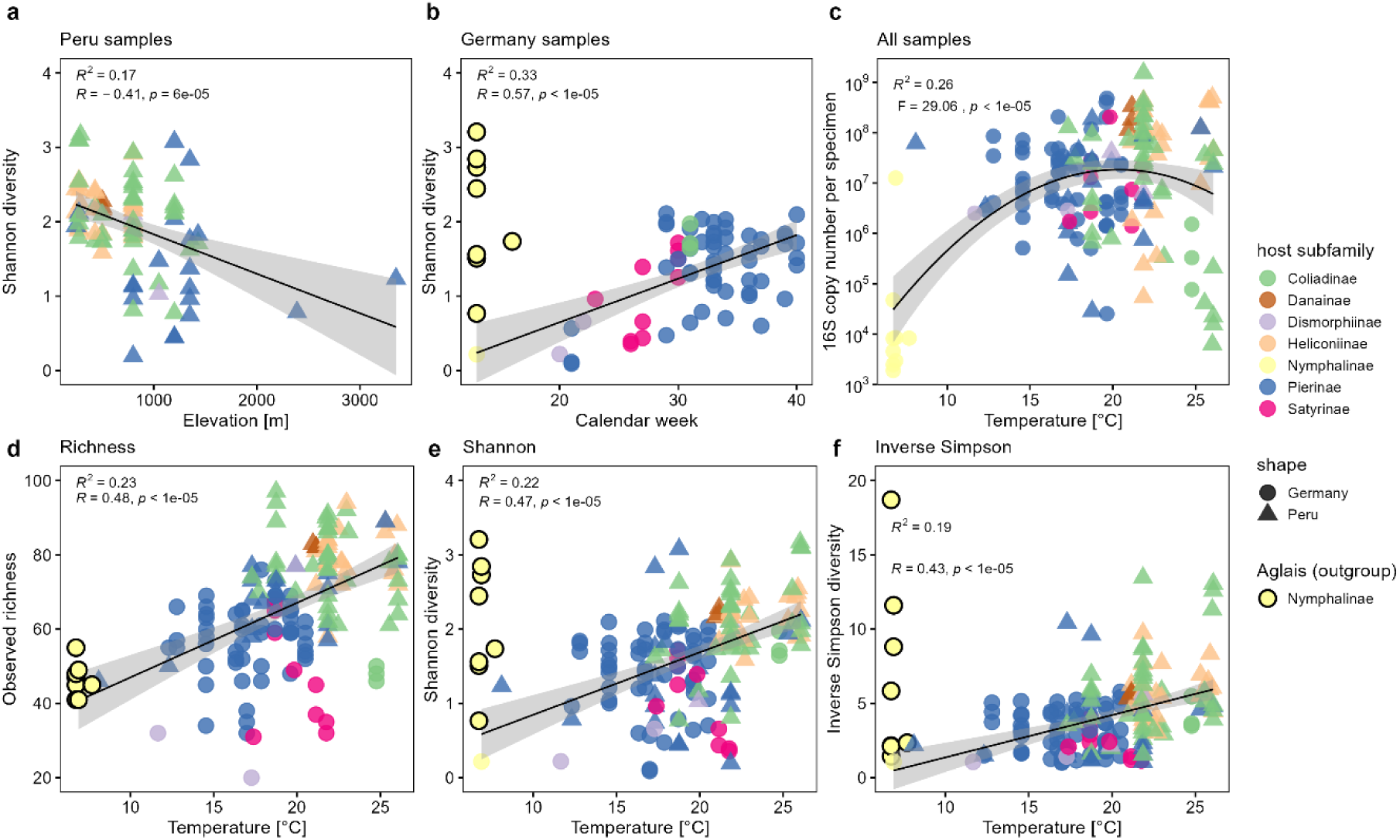
Influence of environmental gradients on adult butterfly microbiota diversity. (**a**) Alpha diversity (only Peru samples) shows linear decrease with elevation (meters above sea level). (**b**) Alpha diversity (only Germany samples) shows linear increase by calendar week. *Aglais* were removed from diversity analyses, due to their low microbial abundance, but illustrated as outgroup within the figure. (**c**) Temperature did not significantly affect bacterial load after accounting for host subfamily. (**d**) For observed microbial richness, both host subfamily and temperature were significant, with host subfamily having the stronger effect (partial R² = 0.27, *F*_6,151_ = 9.42, *p* < 10^-8^), while temperature had a smaller but significant effect (partial R² = 0.03, *F*_1,151_ = 4.63, *p* = 0.033). (**e**) For Shannon diversity, the effect was attributable to host subfamily, whereas temperature was not significant. (**f**) For inverse Simpson diversity, host subfamily had the stronger effect (partial R² = 0.14, *F*_6,151_ = 3.97, *p* = 0.001), while temperature was still significant (partial R² = 0.03, *F*_1,151_ = 4.32, *p* = 0.039).

**Supplementary Figure S3.**
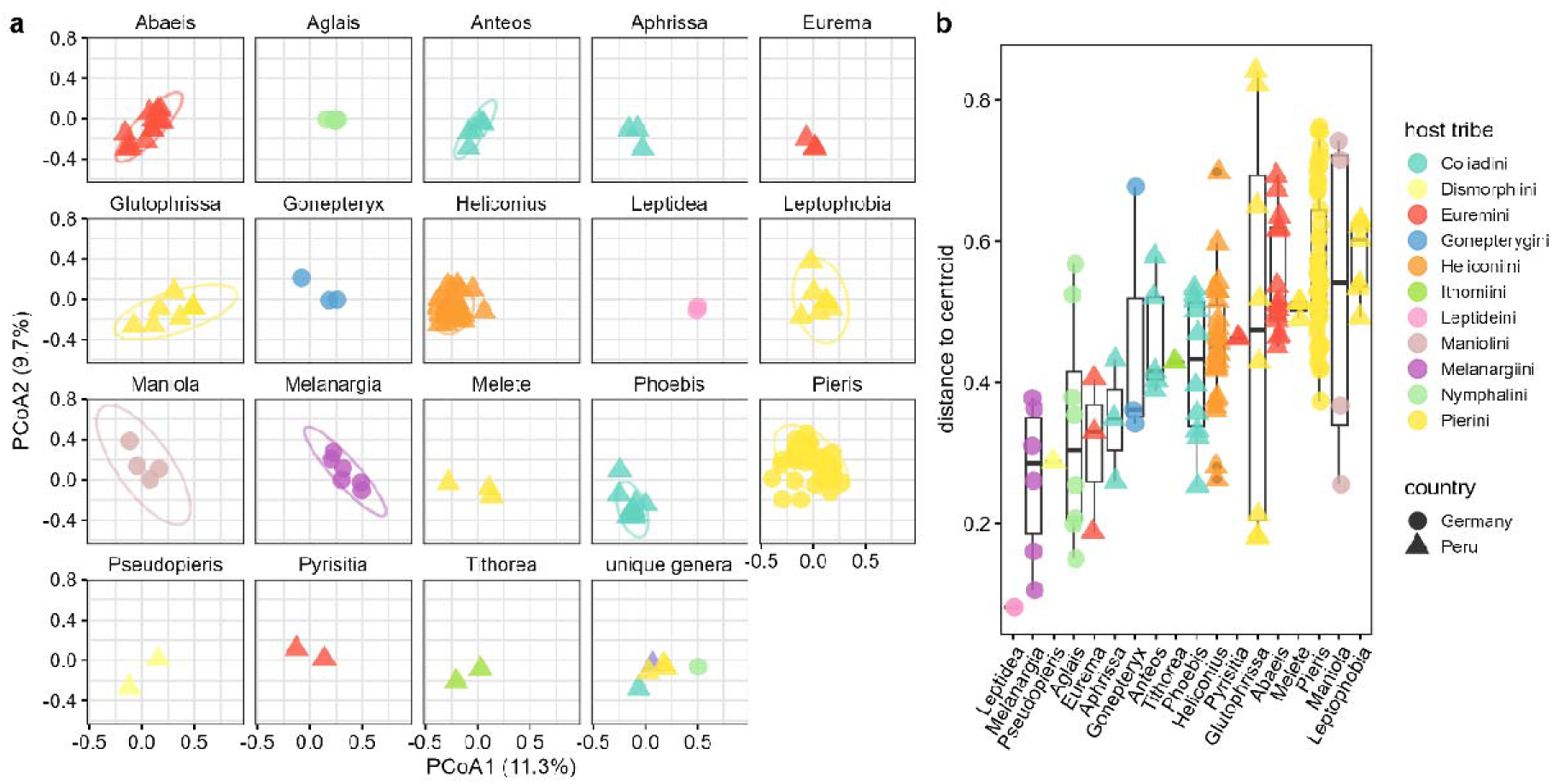
Adult butterfly microbiome dissimilarity and dispersion across host genera. (**a**) Beta diversity of the adult butterfly microbiota using PCoA based on Bray-Curtis distances. Samples are colored by host tribe and panels faceted by host genus (unique genera with less than two replicates were excluded). (**b**) Microbiota of adult butterfly genera differs in beta dispersion (colored by host tribe).

**Supplementary Figure S4.**
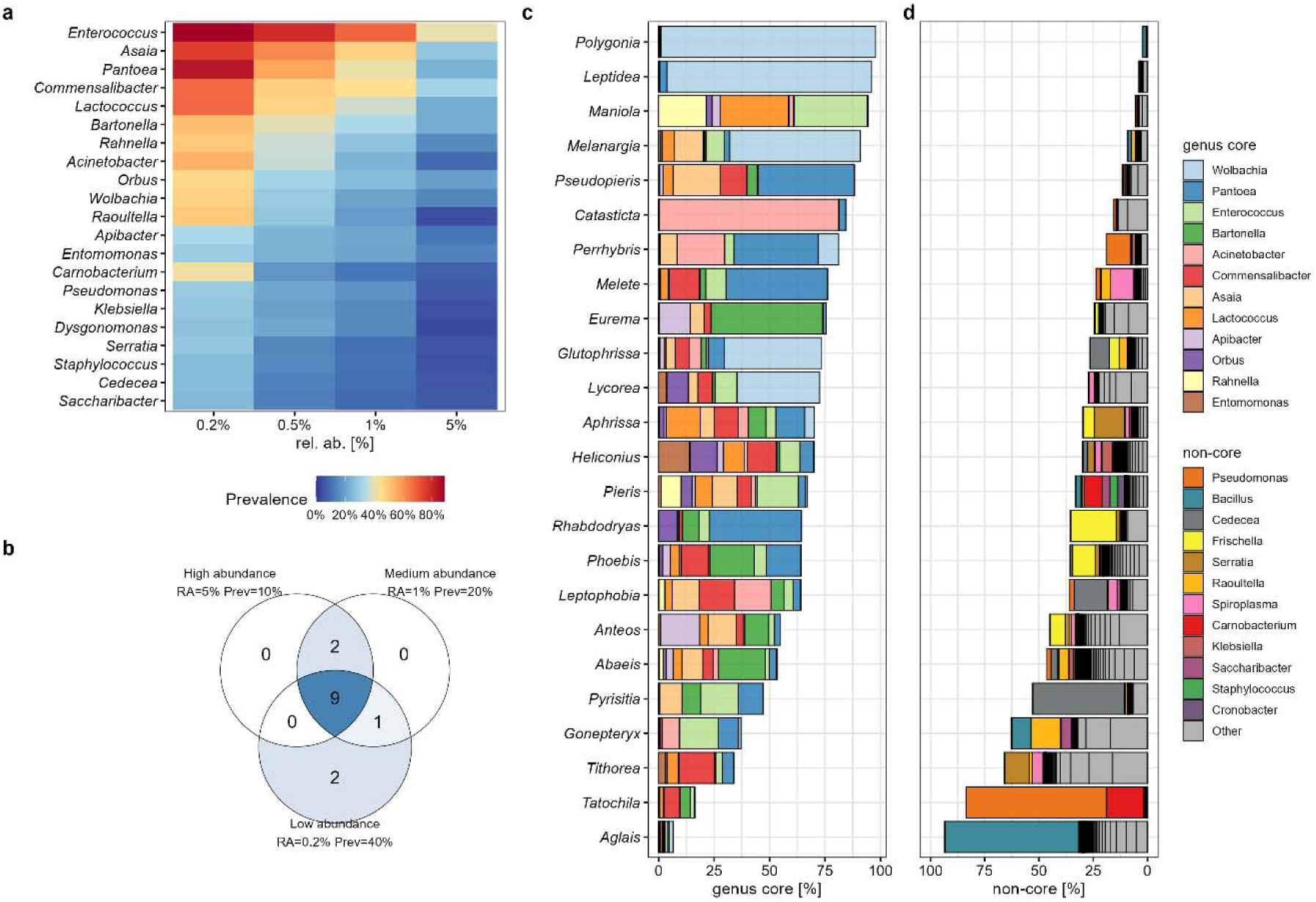
Selection of the common genus core microbiota. (**a**) Heatmap highlighting the most abundant and prevalent genera among all butterfly samples. (**b**) Venn diagram shows overlap of common core genera when applying different core definitions. High abundance definition (5% rel. abundance 10% prevalence) results in 11 core genera (missing *Acinetobacter*); Medium abundance definition (1% rel. abundance 20% prevalence) results in 12 core genera (as used in this study). Low abundance definition (0.2% rel. abundance 40% prevalence) results in 12 core genera replacing *Entomomonas* & *Apibacter* with *Carnobacterium* & *Raoultella*. (**c**) Average relative abundance of the 12 common genus core taxa among individual butterfly genera. (**d**) Remaining non-core genera belonged to the same bacterial groups e.g. LAB (*Carnobacterium*), enteric bacteria (*Cedecea*, *Serratia*, *Roultella*), or Orbacea (*Frischella*).

**Supplementary Figure S5.**
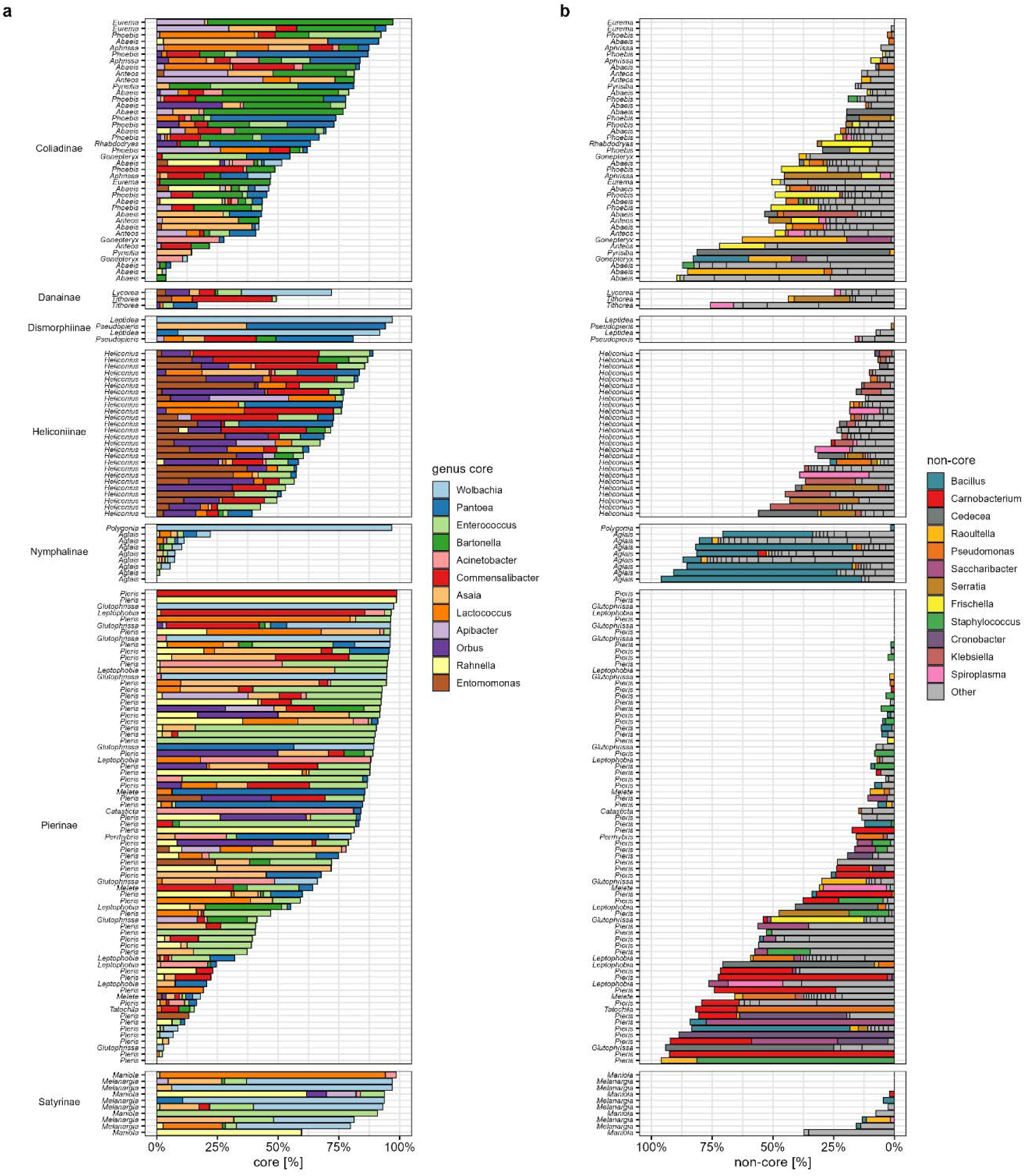
Adult butterfly microbiome composition across individual samples. (**a**) Average relative abundance of the 12 common genus core taxa per individual sample grouped by host subfamily. (**b**) Remaining non-core genera per sample grouped by host subfamily. Relative abundances < 1% are not shown.

**Supplementary Figure S6.**
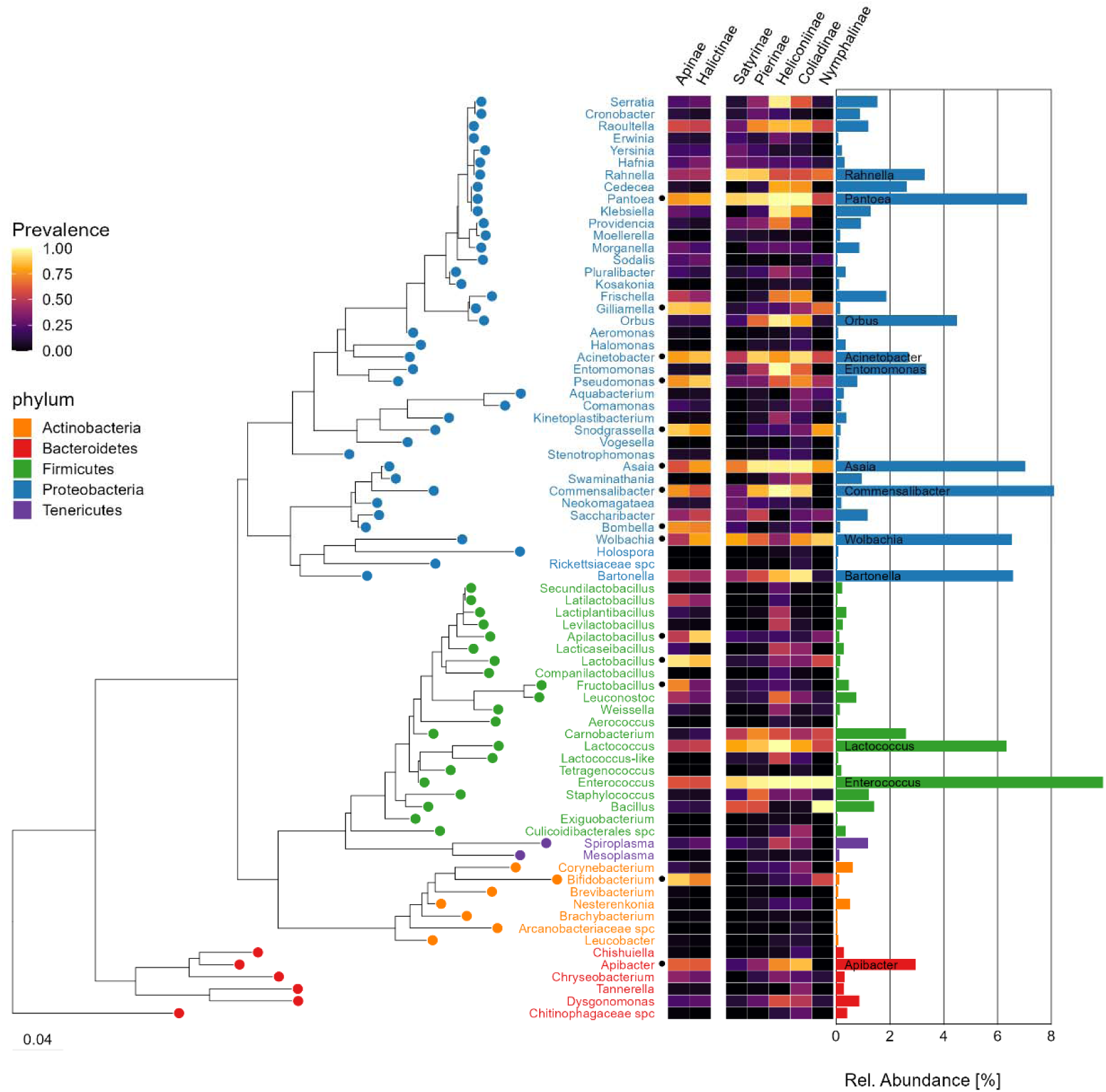
Full genus-level annotation, prevalence and abundance of the adult butterfly microbiota. Phylogenetic grouping (SINA RAxML tree) and relative abundance of the 80 most abundant bacterial genera from adult butterflies. Expansion of Figure 4 including all bacterial genus names. Prevalence in wild bee microbiomes from sympatric Apinae and Halictinae is shown for comparison. Black dots indicate common core genera identified from the bee dataset.

**Supplementary Figure S7.**
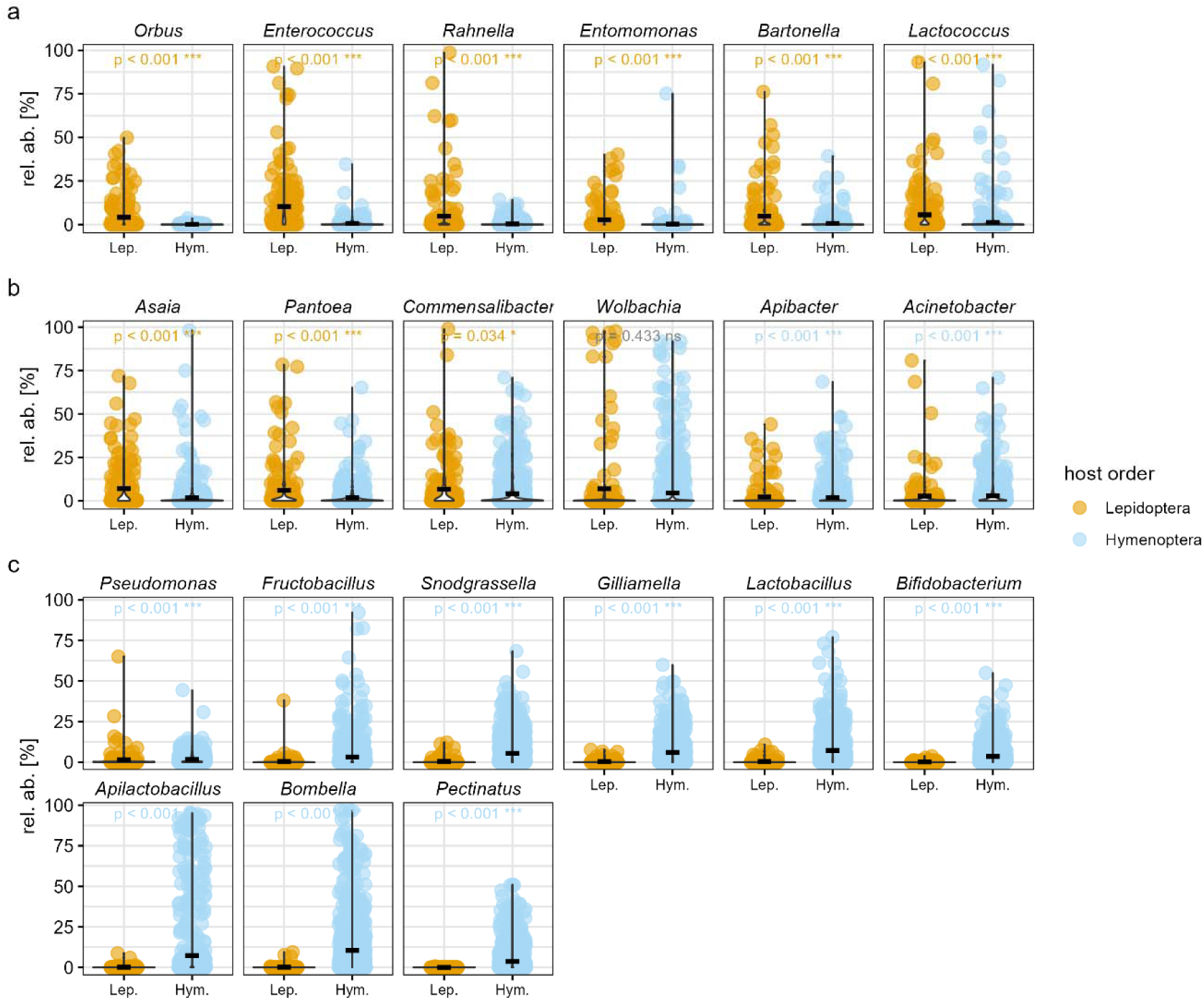
Comparison of the common genus core microbiota of Lepidoptera and Hymenoptera. (**a**) Six genera are unique to the common core of Lepidoptera. (**b**) Six genera show taxonomic overlap between the common cores of Lepidoptera and Hymenoptera. (**c**) Nine genera are unique to the common core of Hymenoptera. Using the same operational common core definition for Lepidoptera and Hymenoptera should facilitate comparability between datasets but does not necessarily represent an optimized threshold for the Hymenoptera dataset, nor is it intended to specifically identify socially transmitted taxa. Wilcoxon rank-sum tests with Benjamini–Hochberg correction for multiple comparisons (Lepidoptera *n* = 167, Hymenoptera *n* = 872).

**Supplementary Figure S8.**
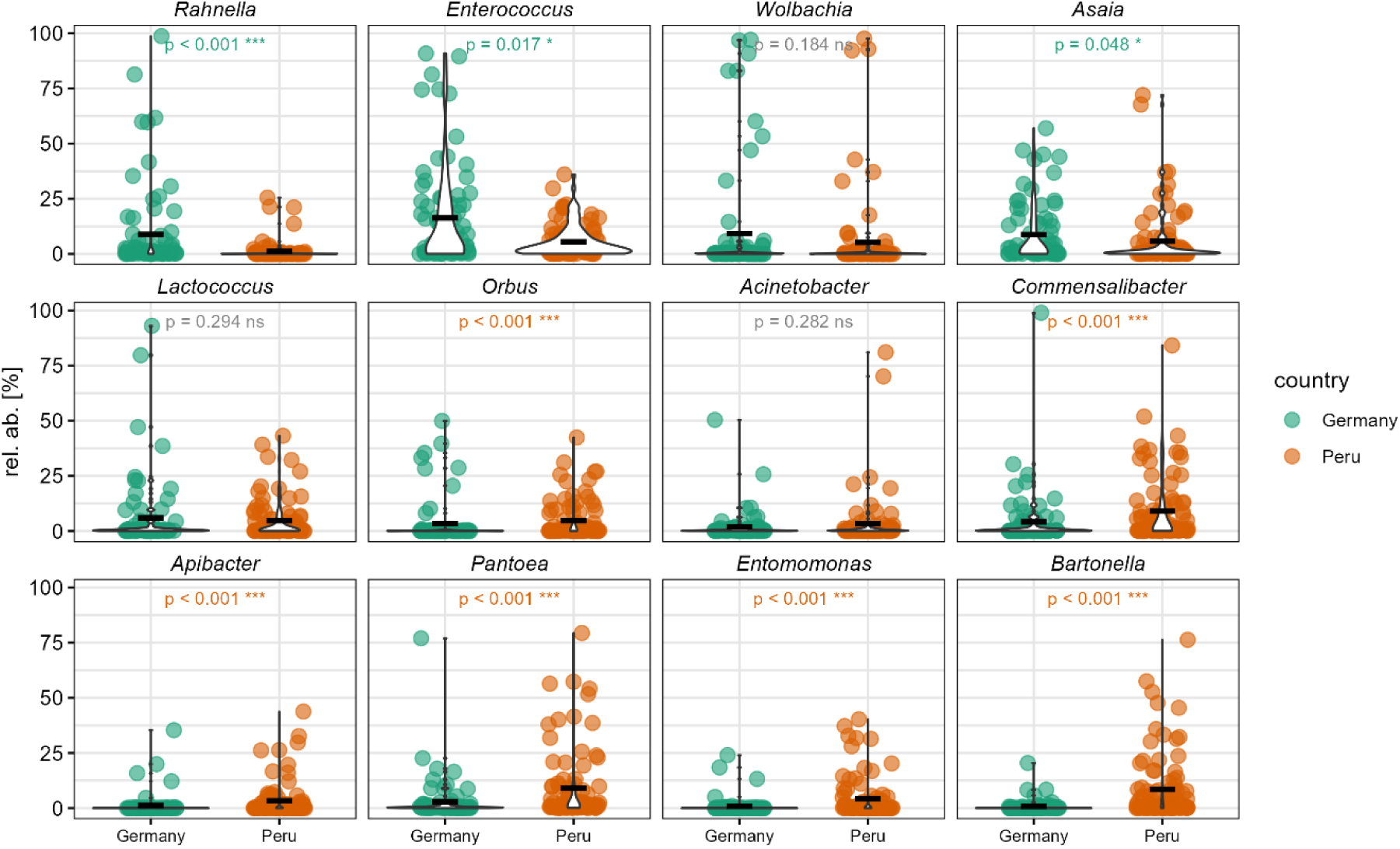
Country-specific distribution of common genus core taxa. Relative abundance of individual core genera among countries and results of Wilcoxon rank-sum Tests following Benjamini–Hochberg correction accounting for multiple comparisons (Germany n = 76, Peru n = 91).

**Supplementary Figure S9.**
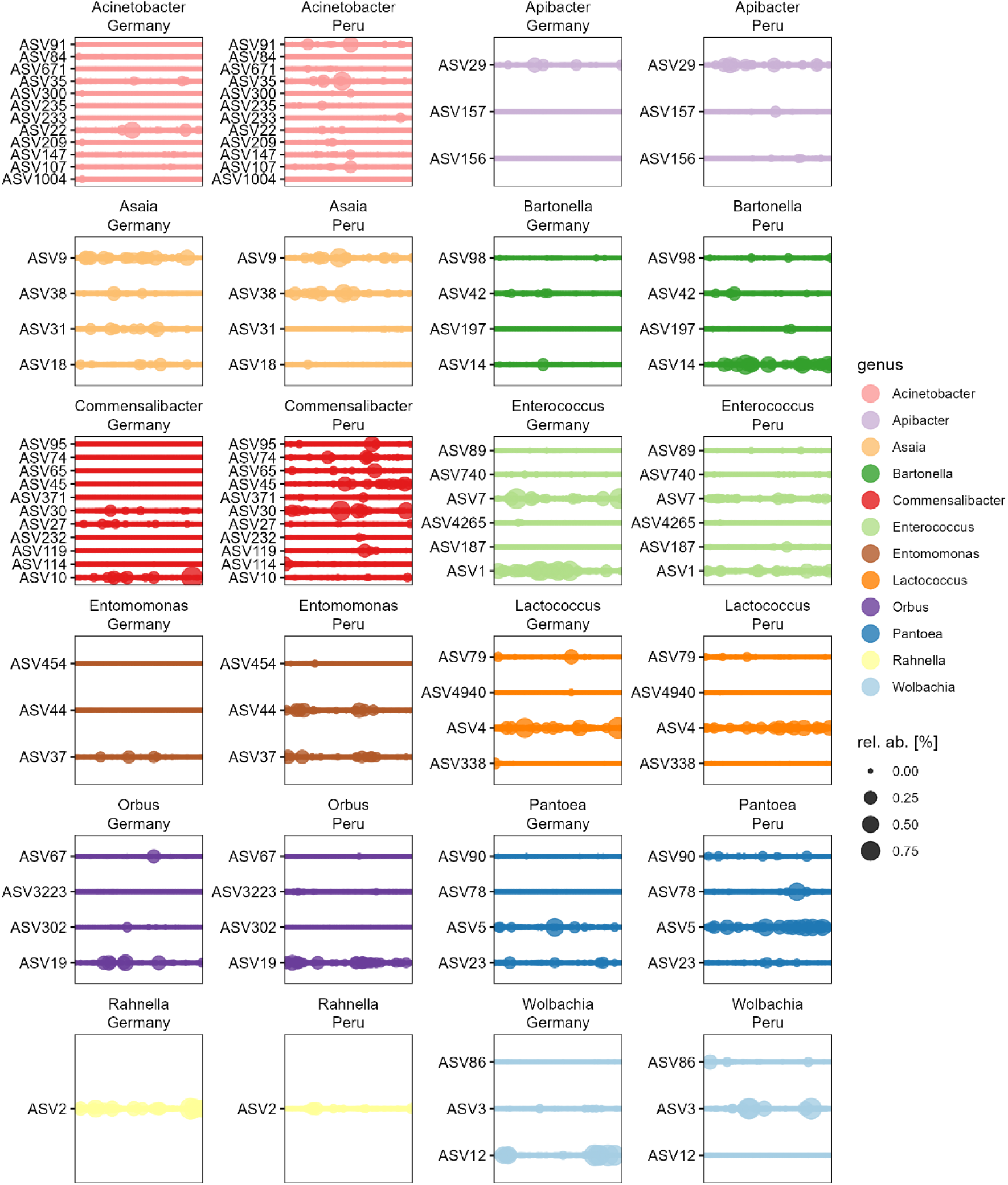
Distribution of common genus core ASVs across countries. Relative abundance and distribution of individual ASVs of the 12 common genus core taxa. Only ASVs with ≥ 2% relative abundance in at least one sample were shown.

**Supplementary Figure S10.**
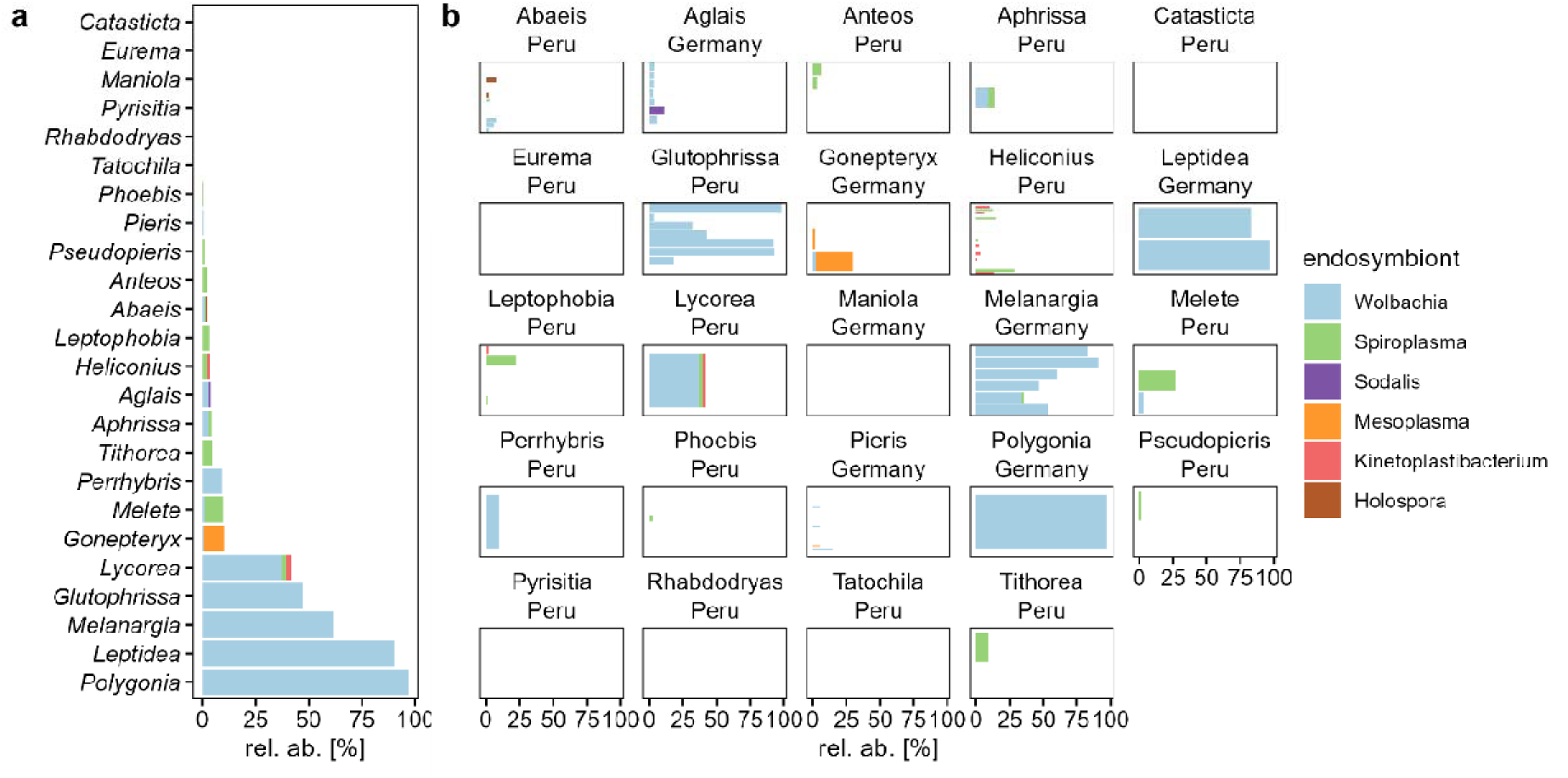
Endosymbionts in the adult butterfly microbiota. (**a**) Average relative abundance of endosymbionts grouped per butterfly genus. (**b**) Relative abundance of endosymbionts among individual samples grouped by genus and country. Relative abundances < 1% are not shown.

**Supplementary Figure S11.**
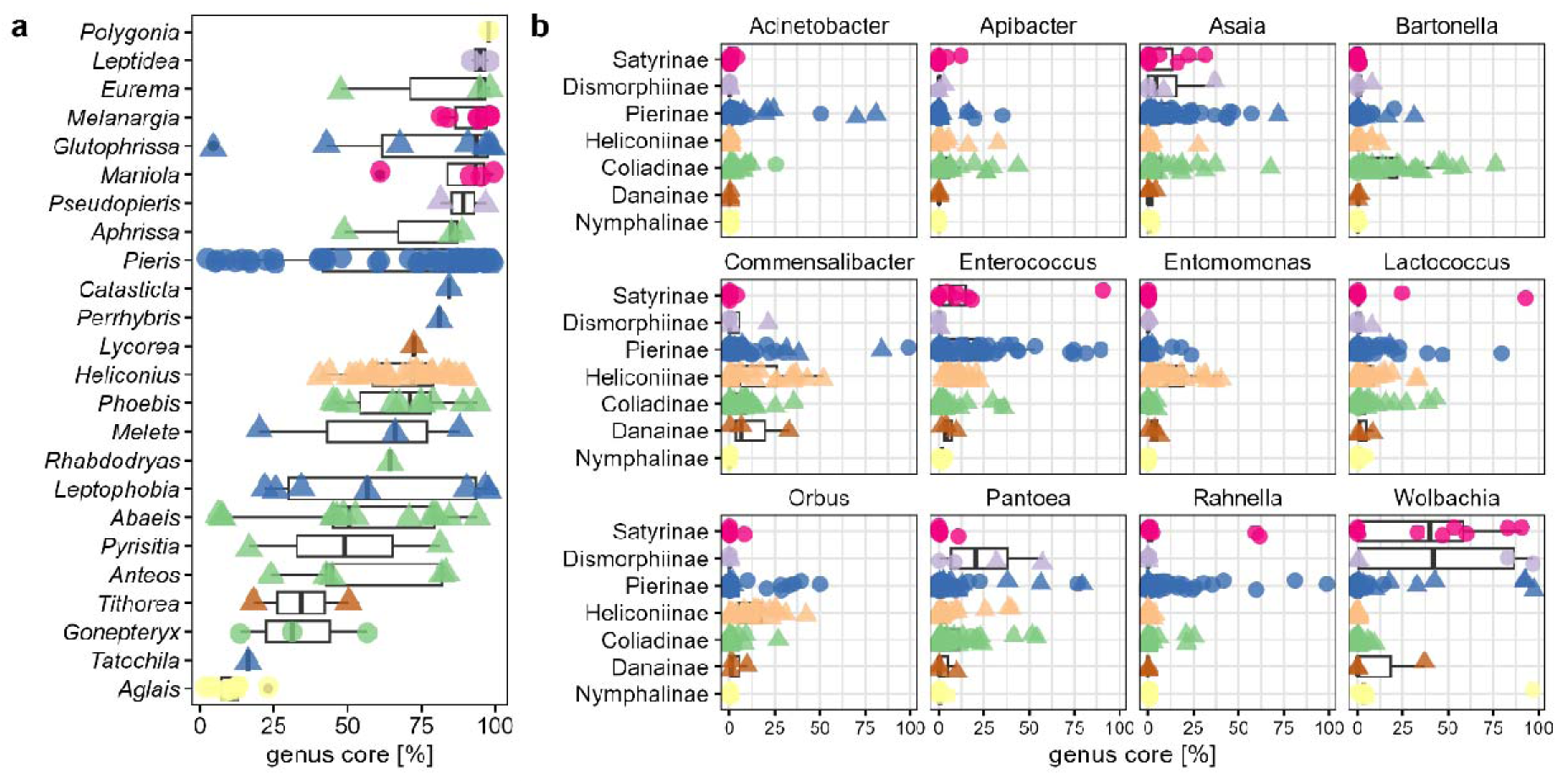
Distribution of common genus core taxa among butterfly samples. (**a**) Total genus core abundance among individual butterfly genera. (**b**) Distribution of individual genus core taxa across butterfly subfamilies. Symbols depict different sampling countries (Circles for Germany, triangles for Peru).

**File: Suppl_table_sample_filter_metadata.csv**

**Supplementary Table S1. Filtering steps and sample metadata**

Removed reads per individual sample (total and percentage) including all metadata and individual BioProject accession numbers.

